# Novel xylan degrading enzymes from polysaccharide utilizing loci of *Prevotella copri* DSM18205

**DOI:** 10.1101/2020.12.10.419226

**Authors:** Javier A. Linares-Pastén, Johan Sebastian Hero, José Horacio Pisa, Cristina Teixeira, Margareta Nyman, Patrick Adlercreutz, M. Alejandra Martinez, Eva Nordberg Karlsson

## Abstract

*Prevotella copri* DSM18205 is a bacterium, classified under Bacteroidetes that can be found in the human gastrointestinal tract (GIT). The role of *P. copri* in the GIT is unclear, and elevated numbers of the microbe have been reported both in dietary fiber-induced improvement in glucose metabolism but also in conjunction with certain inflammatory conditions. These findings raised our interest in investigating the possibility of *P. copri* to grow on xylan, and identify the enzyme systems playing a role in digestion of xylan-based dietary fibers in *P. copri*, which currently are unexplored. Two xylan degrading polysaccharide utilizing loci (PUL10 and 15) were found in the genome, with three and eight GH-encoding genes, respectively. Three of the eight gene products were successfully produced in *Escherichia coli*: One monomeric two-domain extracellular enzyme from GH43 (subfamily 12, in PUL10, 60 kDa) and two dimeric single module enzymes from PUL15, one extracellular GH10 (41 kDa), and one intracellular GH43 subfamily 1 enzyme (37 kDa). The GH43_12 enzyme was hydrolysing arabinofuranose residues from different substrates, and a model of the 3D-structure revealed a single arabinose binding pocket. The GH10 (1) and GH43_1 are cleaving the xylan backbone. Hydrolysis products of GH10 (1) were DP2-4, and seven subsites (−3 to +4) were predicted in the 3D-model of the GH10 active site. GH43_1 mainly produced xylose (in line with its intracellular location). Based on our results we propose that in PUL15, GH10 (1) is an extracellular endo-1,4-β-xylanase, that hydrolyses mainly glucuronosylated xylan polymers to xylooligosaccharides (XOS); while, GH43_1 in the same PUL, is an intracellular β-xylosidase, catalysing complete hydrolysis of the XOS to xylose. In PUL10, the characterized GH43_12 is an arabinofuranosidase, with a role in degradation of arabinoxylan, catalysing removal of arabinose-residues on xylan polymers.

## Introduction

*Prevotella copri* is a Gram-negative non-spore forming anaerobic bacterium, classified under the phylum Bacteroidetes. *P. copri* can be found in the human gastrointestinal tract (GIT) and has been isolated both from human faeces and human oral cavities (Hayashi et al. 2007).

The microbiota of the human gastrointestinal tract (GIT) plays an important role in human health but is a complex system with substantial individual variation in composition and in response to diet. The species distribution, diversity and metabolic outputs of the gut microbiota affects the host in a way that can be either beneficial or harmful. Some microbial species in the GIT play important roles in human health such as providing nutrients, protecting against pathogens, modulating the host metabolism and immune system by secretion of metabolites, while others may have harmful effects such as increased risk of inflammation and disease. The role of *P. copri* in the GIT is still not clear. *P. copri* has been reported as an organism found in elevated numbers in patients with rheumatoid arthritis, leading to a reduction in the abundance of other bacterial groups (Scher et al. 2013). *P. copri* has, however, also been reported as a potential beneficial microorganism leading to dietary fiber-induced improvement in glucose metabolism in certain individuals (Kovatcheva-Datchary et al. 2015).

Dietary fiber consists of polysaccharides and oligosaccharides, and is the portion of plant-derived food that cannot be completely broken down by human digestive enzymes. The polysaccharides found in the fibers include: cellulose and other β-glucans, xylans, pectins, and resistant starch. Many of these dietary fibers can be utilized by the gut microbiota. In consumption trials using barley kernel-based bread, metagenomic analysis of healthy responders showed that the gut microbiota was enriched in *P. copri* and had increased potential to ferment complex polysaccharides (Kovatcheva-Datchary et al. 2015). This finding raised our interest in investigating *P. copri* and its enzyme systems, as well as the possibility of *P. copri* to grow on xylan, and/or xylooligosaccharides being emerging prebiotics. The human genome does not contain genes for xylanases, xylosidases, or arabinofuranosidases and experimental studies have confirmed that XOSs and arabinoxylooligosaccharides (AXOSs) are not degraded by human saliva, artificial gastric juice, pancreatin, or intestinal mucosa homogenate, while specific microorganisms are able to grow on XOS/AXOS allowing selective stimulation of these groups in the GIT (Nordberg Karlsson et al. 2018).

Degradation of dietary fibers often requires concerted action of several carbohydrate active enzymes, and in Bacteroidetes, bacterial species growing in competitive environments (e.g. *P. copri* in the GI-tract) have organized genes encoding various carbohydrate active enzymes, proteins and transporters required for saccharification of complex carbohydrates in colocalized clusters called polysaccharide utilization loci (PULs), which are strictly regulated (Grondin et al. 2017).

The presence of PULs is a unique feature of Bacteroidetes genomes, and is a term used to describe clusters of colocalized, coregulated genes, the products of which orchestrate the detection, sequestration, enzymatic digestion, and transport of complex carbohydrates (Bjursell, Martens, and Gordon 2006).

PULs encode a number of complementing cell surface glycan-binding proteins (SGBPs), TonB-dependent transporters (TBDTs) and CAZymes, mostly Glycoside Hydrolase (GHs), but also Polysaccharide Lyases (PLs) and Carbohydrate Esterases (CEs), and complex carbohydrate sensors/transcriptional regulators (Hamaker and Tuncil 2014; Martens et al. 2014). The complexity of PULs often increases with the complexity of their cognate substrates and may dependent on the substrate also include proteases, sulfatases, and phosphatases (Abbott et al. 2015; Barbeyron et al. 2016; Renzi et al. 2015).

The focus of this work was to identify and characterize enzyme candidates that play a role in the degradation of xylan based dietary fibers in *P. copri*. In cereal grains, neutral arabinoxylans (AX) with xylopentaose (Xylp) residues substituted at position 3 and/or at both positions 2 and 3 of Xylp by α-L-arabinofuranoside (Araf) units, representing the main xylan components (Ebringerová and Heinze 2000). The xylan polymer can be hydrolysed by xylanases that are produced by a range of different microorganisms.

Most of the main-chain acting endo-xylanases (EC 3.2.1.8) known to date have evolved from two main scaffolds: the TIM-barrel (α/β)_8_ (found in the glycoside hydrolase (GH) families: GH5, GH10 and GH30, and the β-jelly roll (found in GH11) (Nordberg Karlsson et al. 2018). In addition, a number of GH families show beta-xylosidase activity (EC 3.2.1.37), and successively remove D-xylose residues from the non-reducing termini. The majority of these enzymes from bacteria are found in GH39 (retaining), and GH43 (inverting).

To gain more details on the activity of xylan degrading enzymes in *P. copri*, the PULs of the bacterium were analysed and candidates potentially involved in xylan degradation were cloned and produced. This led to the identification and characterization of two xylan degrading PULs, and the characterization of three novel enzymes from *P. copri* PULs.

## Materials and methods

### Preparation of arabinoxylan (AX) and arabinoxylan-oligosaccharide (AXOS) extracts from Brewer’s spent grain as carbon source for utilization trials in Prevotella copri

Brewer’s spent grain (BSG), provided by Viking Malt, was prepared by mashing Pilsner malt at temperatures between 45-70°C (1°C min^−1^). After the wort was cooled (~20°C) and filtered the solids were dried at 48-55°C (1°C min^−1^) for 20h. From the BSG, four xylan fractions were prepared for screening of carbon source utilization in *P. copri*: Water-extractable arabinoxylan (WE-AX), alkaline-extractable arabinoxylan (AE-AX), pellet arabinoxylan (Pellet-AX) and a combination of the three xylan fractions (WAP-AX). (**Table 1**)

**Table 1.**
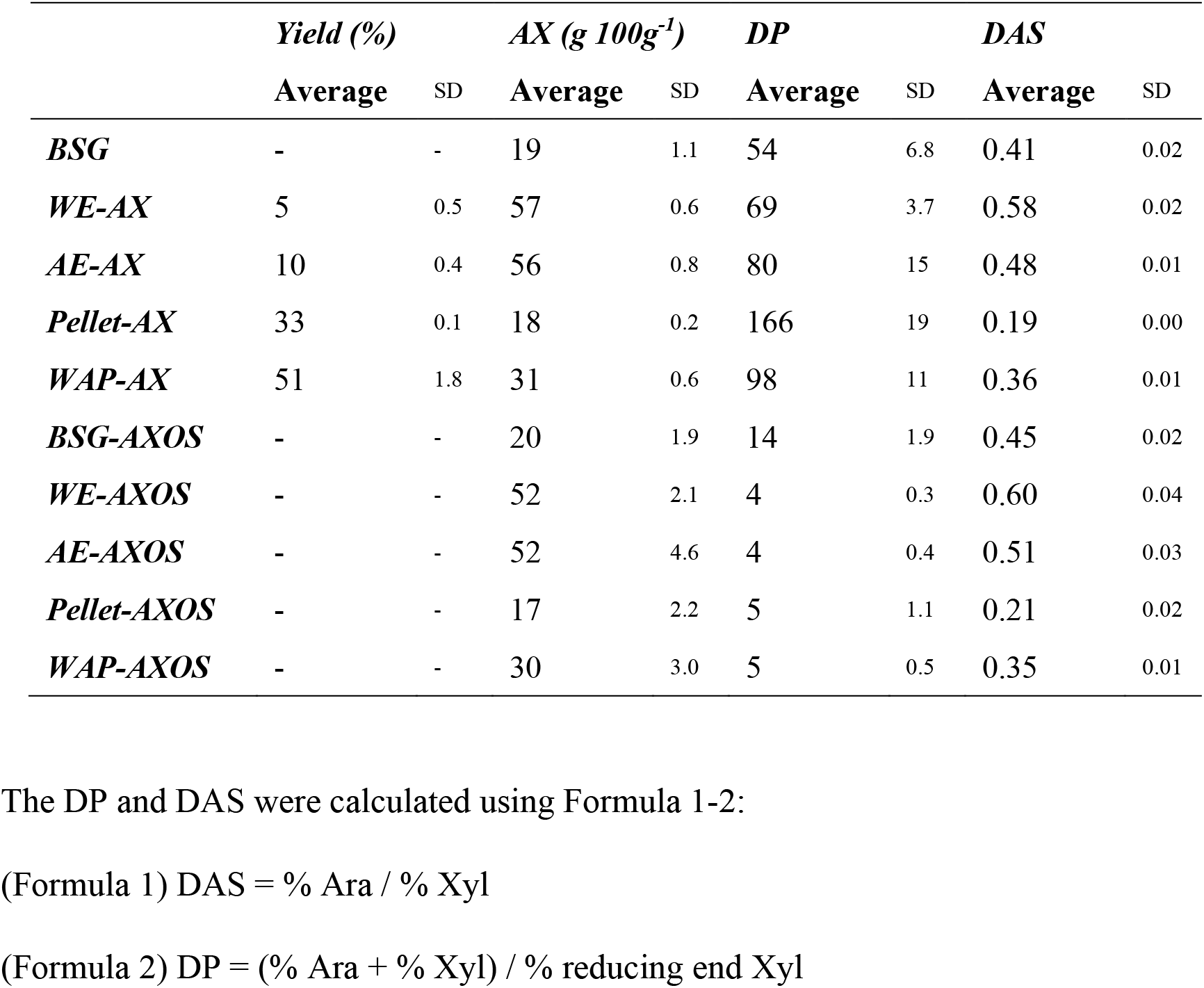
Arabinoxylan (AX) content, degree of polymerization (DP) and degree of arabinose substitution (DAS) in fractions of AX and arabinoxylooligosaccharide (AXOS) prepared as carbon source for utilization trials in *Prevotella copri*.

WE-AX and AE-AX were prepared as described previously (Sajib et al. 2018). BSG (5 g, dry weight) was destarched by incubation with amylase (0.2 mL, Termamyl 120 L, alpha-amylase from *Bacillus licheniformis* – Type XII-A, Sigma-Aldrich) in sodium phosphate buffer (125 mL, 20 mM, pH 6.9) in a water-bath at 90°C for 90 min. After centrifugation the pellet was first washed with 125 mL and then resuspended in 50 mL ultrapure water. The suspension was autoclaved at 121°C for 15 h followed by centrifugation to recover the WE-AX containing supernatant. WE-AX was then precipitated with four volumes of 99.5% ethanol, kept overnight at 4°C and washed four times with 80% etanol before drying. To obtain AE-AX, the pellet of the autoclaved suspension was treated with KOH (25 mL, 0.5M) for 2 h at 40°C in a shaking water-bath, centrifuged, and the supernatant saved. The pellet was washed (25 mL, water) and centrifuged, and the supernatant was combined with the one previously collected The supernatants were, neutralized with HCl and precipitated with four volumes of 99.5% ethanol, kept overnight at 4°C and the precipitated AE-AX was washed four times with 80% ethanol before drying. The Pellet-AX was obtained from the pellet remaining after the alkaline treatment and wash. The combined arabinoxylan extract (WAP-AX) was prepared in the same way as WE-AX, AE-AX, and Pellet-AX: After amylase and hydrothermal treatment, the pellet was treated with KOH as for AE, neutralized and all the suspensions and pellets were merged, neutralized and precipitated with ethanol.

A part of the BSG and each of the four extracts were enzymatically treated with endo-xylanase (Pentopan 500 BG, EC 253.439.7, Novozymes), resulting in five oligosaccharide fractions: BSG arabinoxylan-oligosaccharides (BSG-AXOS) Water-extractable arabinoxylan-oligosaccharides (WE-AXOS), alkaline-extractable arabinoxylan-oligosaccharides (AE-AXOS), pellet arabinoxylan-oligosaccharides (Pellet-AXOS) and AXOS from the combined extracts (WAP-AXOS). Suspensions were prepared with extract or BSG in sodium phosphate buffer (20 mM), and the xylanase was added to an equivalent of 1U/g AX and incubated at 40°C for 5 h. The reaction was stopped by boiling, 5 min, and kept frozen until freeze-dried.

The arabinoxylan content, average degree of polymerization (DP) and degree of arabinose substitution (DAS) was determined in all AX and AXOS fractions, including BSG (**Table 1**).

Arabinose and xylose content and xylan reducing ends were determined by a gas chromatographic (GC) methodology (Theander et al. 1995) that comprises hydrolysis, reduction and acetylation of the saccharides. Briefly, the samples were hydrolysed with 12M H_2_SO_4_, and incubated for 1h at 30°C followed by autoclaving (1h, 121°C). Reduction was done by adding NH_3_ and KBH_4_ and incubating at 40°C for 1 h. The reaction was stopped with CH_3_COOH. The samples were then acetylated by adding 1-methylimidazol, acetic anhydride, 99.5% ethanol, water and 7.5M KOH. The organic layer containing the acetylated hydrolysed monosaccharides were dried with Na_2_SO_4_ and quantified by GC. For analysis of xylan reducing ends, the reduction reaction was performed prior to hydrolysis and the acetylation was similar to a method adapted from Courtin (Courtin, Van den Broeck, and Delcour 2000). A few drops of octanol (~50 μl) were added to the reducing reaction to avoid foaming when adding CH_3_COOH.

### Growth screening and analysis of cultivation broth

To screen growth of *Prevotella copri* on AX and AXOS substrates, a Peptone Yeast Glucose (PYG) medium was prepared according to the guidelines by DSMZ (Deutsche Sammlung von Mikroorganismen und Zellkulturen, Germany, Medium 104) without adding the sugar carbon source (glucose). The medium was distributed into serum flasks, under anaerobic conditions, and then either BSG, an extract or enzymatically treated extracts was added to a final concentration of 25 mg mL^−1^. The flasks were rubber sealed and autoclaved (15 min, 121°C). After cooling (<40°C), horse serum (5% v/v) was added to the medium.

The flasks with the respective medium were pre-heated at 37°C, approximately 30 min prior to inoculation. Each flask was inoculated with 1 mL *P. copri* inoculum/20 mL medium. The pre-inoculum consisted of an exponential phase *P. copri* culture in PYG medium (with glucose as carbon source).

The pH of the cultures was analysed 48 and 72h after inoculation, and at these times samples were also withdrawn for analysis of DAS and utilization of xylans and AXOS using the GC-methodology described above. All procedures were done in duplicate.

### Gene synthesis and cloning

Sequences of putative CAZymes coding genes from *Prevotella copri* DSM 18205 were retrieved from the Polysaccharide-Utilization Loci DataBase (Terrapon et al. 2018). GH10 and GH43 encoding sequences were selected from two clusters and then synthesized and cloned into the expression plasmid pET-21b(+) through GenScript® services (Piscataway, NJ, USA).

Prior to cloning, the deduced amino acid sequences were analysed for the presence of a signal peptide (Petersen et al. 2011), and the genes were subsequently synthesized and cloned including native signal peptides. The recombinant plasmids obtained were dissolved in Milli-Q water (100 μmol ml^−1^), and finally 10 ng were transferred to *E. coli* strain BL21 (DE3) competent cells by thermal shock transformation.

### Protein expression and purification

The recombinant proteins were expressed in *E. coli* BL21(DE3). The cells grew at 37°C or at 25°C (LB medium) until reaching an optical density (OD) at 600 nm of 0.6. At this stage, expression was induced by adding 1.0 or 0.7 mM IPTG (isopropyl b-D-1-thiogalactopyranoside, as specified in Table 1), and expression in the respective culture continued for 4 h at 37°C or 24 h at 25°C (LB medium). The cultures were then harvested, and cell pellets were collected by centrifugation, washed, and resuspended in binding buffer (BB, sodium phosphate 20 mM, NaCl 500 mM, pH 7.4), followed by sonication as described (Faryar et al. 2015).

The cell extracts obtained were centrifuged at 23,000 × *g*, 4 °C, 20 min, and each His-tagged enzyme was purified by immobilized metal ion affinity chromatography (IMAC) using a HiTrap FF affinity 5 mL column (GE Health Care, Germany) on an ÄKTA start system (Amersham Biosciences, Uppsala, Sweden). Bound proteins were eluted using an elution buffer (EB, sodium phosphate 20 mM, NaCl 500mM, imidazole 500 mM, pH 7.4), and finally dialyzed against BB overnight to remove the imidazole. Protein purity was evaluated by SDS/PAGE in a 4-20% precast polyacrylamide gradient gel (BioRad, Copenhagen, Denmark). Protein concentrations were estimated according to Bradford, (Bradford 1976), using a reagent from BioRad (Copenhagen, Denmark).

### Molecular mass determination of purified enzymes

The molecular mass for each expressed enzyme was estimated by size exclusion chromatography as described by Michlmayr et al. (Michlmayr et al. 2013), using a HiPrep™ 16/60 Sephacryl® S-300 HR column (120 mL; GE Healthcare) at a flow rate of 0.5 ml min^−1^, with sodium phosphate 50 mM, NaCl 150 mM, pH 7.0. Calibration using molecular weight standards in the range 20–700 kDa (Sigma-Aldrich, Saint-Louis, MO, USA), allowed estimation of the theoretical molecular mass from Compute pI/MW, ExPASy (https://web.expasy.org/compute_pi/).

### Enzyme activity for synthetic and natural substrates

Reaction mixtures using *p*-nitrophenyl derivatives as synthetic substrates (1 mM) were performed in 50 mM citrate-phosphate buffer (CPB) pH 5.5 at 37 °C, and 0.1 mg mL^−1^ of each enzyme. The following compounds were purchased from Megazyme (Chicago, USA): *p*-nitrophenyl-α-L-arabinofuranoside (*p*-NPAra), *p*-Nitrophenyl-β-D-xylopyranoside (*p*-NPXyl1), *p*-nitrophenyl-β-xylobioside (*p*-NPXyl2), *p*-nitrophenyl-β-xylotrioside (*p*-NPXyl3), *p*-nitrophenyl-β-D-glucopyranoside (*p*-NPGlu), *p*-nitrophenyl-β-D-galactopyranoside (*p*-NPGal), and *p*-nitrophenyl-β-D-mannopyranoside (*p*-NPMan). The absorbance of the *p*-nitrophenol (*p*-NP) produced was measured continuously for 5 min at 400 nm in a microplate reader (Multiscan GO, Thermo Scientific). Enzyme activity was expressed in Units (U) and was defined as μmol of *p*-NP released per min.

Xylans from birchwood and beechwood were purchased from Sigma (USA); rye flour arabinoxylan (soluble), debranched arabinan and sugar beet arabinan were from Megazyme (Ireland). Quinoa xylan was prepared according to Gil-Ramirez (Gil-Ramirez et al. 2018). The enzymatic reaction mixtures, consisting of 360 μl substrate (1% w/v) and 40 μl enzyme solution (0.1 mg mL^−1^) in CPB 50 mM pH 5.5, were incubated at 37 °C for 5 min. The reactions were stopped by addition of 600 μl of 3,5-dinitrosalicylic acid (DNS) and immediately boiled for 10 min. The released sugars were measured from 250 μl aliquots from the enzymatic assays at 540 nm in a microplate reader (Hero et al. 2018). Enzyme activity was expressed in Units (U) and was defined as μmol of reducing sugars released per min.

All samples were analysed in triplicate and mean values and standard deviations were calculated. The specific activity was determined as U per mg of total protein content in the sample.

### pH and temperature optima

Estimation of optimal pH in the pH-range of pH 4.0 to 7.5 was assessed at 37 °C using *p*-NPAra (1 mM) as substrate in 50 mM CPB. For each pH value, a calibration curve was plotted with *p*-NP as standard (Merck, Darmstadt, Germany). Endo-xylanase activity was determined using beechwod xylan (1% w/v) prepared in a pH range of 3.0 to 10.0 in 50 mM CPB and 50 mM glycine-sodium hydroxide buffer. The optimal temperature was evaluated in a range of 20 to 70 °C, the same substrates were used at pH 5.5. In all cases, the samples were incubated at 37°C for 5 min and the enzymatic activity was then determined as described above.

### Hydrolysis profile for arabinoxylooligosaccharides (AXOS)

Arabinoxylooligosaccharides A^2^XX, XA^3^XX and A^2,3^XX purchased from Megazyme (Irland), were used as substrates for testing arabinofuranosidase activity of GH43_12. The reactions were prepared with 1 mM substrate, 50 mM CPB buffer pH 5.5 and 50 mg mL^−1^ of enzyme. The mixture was incubated at 37°C for 30 min. Then, the reactions were stopped by heating at 95°C for 5 min. All reactions were performed in triplicates with corresponding negative controls consisting in the same reaction-mixture compositions, except enzyme. The products were dilute in ultrapure water (50 mL sample/450 mL water) and analysed by high performance anion exchange chromatography with amperometric detection (HPAEC-PAD, Dionex, San Diego, CA, USA) with Dionex™ CarboPac™ PA200 (Linares-Pastén et al. 2017). Standards of arabinose, A^2^XX, XA^3^XX and A^2,3^XX, in a concentration range from 1 to 20 mM, were used.

### Hydrolysis profile for xylan-containing substrates

Birchwood xylan, beechwood xylan, rye flour arabinoxylan and quinoa xylan (1% w/v) were used as natural substrates for the respective enzyme (0.1 mg mL^−1^). The assay conditions were 37 °C, pH 5.5 for 5 min, and the reaction was stopped by heating at 95 °C for 5 min. Products were analyzed by HPAEC-PAD, as described above. Standards of D-(+)-xylose (Sigma-Aldrich), 1,4-β-D-xylobiose, 1,4-β-D-xylotriose, 1,4-β-D-xylotetraose, 1,4-β-D-xylopentaose and 1,4-β-D-xylohexaose (Megazyme) were used to identify and quantify the peaks in the chromatograms.

### Kinetic constants

Initial reaction rates for GH43_1 from PUL15 at 0.1 mg mL^−1^ were assayed with xylans and arabinoxylan in the range of 0-25 mM, at pH 5.5 and 37 °C. Samples were taken after 1 and 5 min and the reactions were stopped by heating at 95 °C for 5 min. The hydrolysis products produced were quantified by HPAEC-PAD as described above.

In the case of GH43_12 from PUL10 and GH10 from PUL15, synthetic substrates *p*-NPAra, *p*-NPXyl2 and *p*-NPXyl3 were evaluated in the range of concentrations of 0-50 mM; 0-5 mM and 0-15 mM, respectively. The reaction conditions were the same as those used in the previous assay, and the *p*-NP production was measured continuously for 5 min at 400 nm.

Data obtained were fitted to the Michaelis–Menten model by nonlinear regression using GraphPad Prism7® built in functions.

### Homology modelling

The three-dimensional structural models of the full-length protein were obtained by homology modelling, using YASARA (Krieger and Vriend 2014). The accuracy of the models obtained was supported by the relatively high identity in amino acid sequences, and by using several crystallographic templates to generate hybrid models. The GH10 enzyme from PUL15 was modelled using as templates the following crystallographic structures (PDB codes): 1UQY, 3NIH, 1N82, 5OFJ, 2Q8X, 5OFK, 3MS8 and 3MUI.

The model of GH43_1 (from PUL15) was built based on the crystallographic structures 4MLG, 5A8C, 4NOV and 3C7F. GH43_12 (from PUL10) was modelled using as templates 5JOW, 5JOZ, 2XEI, 1YIF, 2EXK, 2YI7, 2EXH, and 2EXJ. The analysis of the structures was performed using CHIMERA (Pettersen et al. 2004). The validation of the models was assessed based on Z-score, which describes how many standard deviations the model quality is away from the average high-resolution X-ray structure. Thus, higher Z-scores are better while negative Z-scores indicate that the model is worse than a high-resolution X-ray structure (Krieger and Vriend 2014).

### Molecular Docking

Molecular structures of an arabinoxylooligosaccharide XXA^2^XX and a xylotriaose, were built with the Avogadro program (Hanwell et al. 2012). Their geometries were optimized at molecular-mechanics level with the force field MMFF94 and the algorithm Steepest Descend, using 1000 steps and convergence of 10^−7^. XXA^2^XX docked into the active site of the receptor GH43_12, while xylotriaose was docked into the active site of GH43_1. Both receptors were modelled as it is described before. Dockings were performed thorough AutoDock (Morris, Huey, and Olson 2008) implemented in YASARA v19.12.14 software (Krieger and Vriend 2014). The molecular models were analyzed with Chimera (Pettersen et al. 2004).

## Results

### Growth trials of Prevotella copri on arabinoxylans and arabinoxylooligosaccharides

*Prevotella copri* DSM18205 is a species classified under the family Prevotellaceae in the phylum Bacteroidetes that has been found in the human gut (Kovatcheva-Datchary et al. 2015). *P. copri* DSM18205 has, like many other species in the different families of Bacteroidetes (El Kaoutari et al. 2013), capacity to utilize different types of polymeric carbohydrate fibers. Yet, the ability of *P. copri* to grow on xylan, and the enzymes of *P. copri* remain unexplored.

To explore the growth of *P. copri* on xylan polymers and oligosaccharides, a number of AX and AXOS substrates were prepared and used as carbon sources in *P. copri* growth trials (Table 2). The substrates were arabinosylated (within the range 0.2 – 0.6), with a degree of polymerization varying from DP4 in the AXOS to a DP >100 for the Pellet-AX fraction (**Table 1**). Irrespective of the DP and DAS, all inoculated cultures displayed a reduction in pH, indicative of growth of *P. copri*, and that the organism was capable to utilize both polymeric and oligomeric xylans (**Table 2**). AX/AXOS consumption was slow for untreated BSG while extraction and/or xylanase treatment caused an increase in consumption rate. The arabinosylation of the substrates displayed an apparent increase after 48h, that was again reduced after 72h cultivation, indicating that non-substituted regions of xylans were first utilized, but that arabinosylated parts of the xylan were consumed later in the cultivation.

**Table 2.** pH and substrate consumption in the cultivation broth. Data are given as average ± standard deviation of duplicate samples

|  | *pH* |  |  | *DAS* |  |  | *Arabinose consumed (%)* |  |
| --- | --- | --- | --- | --- | --- | --- | --- | --- |
|  | 0h | 48h | 72h | 0h | 48h | 72h | 48h | 72h |
| *BSG* | 7.2±0.07 | 5.0±0.01 | 4.9±0.02 | 0.4±0.02 | 0.5±0.03 | 0.6±0 | 1.8±0.2 | 3.6±0.3 |
| *WE-AX* | 7.4±0.07 | 4.8±0.04 | 4.9±0.01 | 0.6±0.02 | 1.2±0.13 | 1.0±0.3 | 4.7±0.1 | 4.8±0 |
| *AE-AX* | 7.4±0.13 | 6.2±1.4 | 6.1±1.5 | 0.5±0.01 | 0.8±0.3 | 0.8±0.4 | 2.5±2.1 | 1.7±1.8 |
| *Pellet-AX* | 7.4±0.07 | 5.4±0.02 | 5.4±0.08 | 0.2±0 | 0.3±0.02 | 0.2±0 | 4.7±0.2 | 3.2±0.2 |
| *WAP-AX* | 7.3±0.08 | 6.3±0.06 | 6.1±0.22 | 0.4±0.01 | 0.3±0 | 0.4±0 | 3.7±0.1 | 3.8±0 |
| *BSG-AXOS* | 7.0±0.08 | 5.0±0.11 | 4.9±0.06 | 0.4±0.02 | 0.4±0.21 | 0.8±0.2 | 3.7±0.1 | 3.3±0.2 |
| *WE-AXOS* | 7.1±0.06 | 5.0±0.10 | 5.0±0 | 0.6±0.04 | 0.7±0.01 | 0.8±0 | 4.0±1.0 | 4.5±0.2 |
| *AE-AXOS* | 7.1±0.06 | 5.2±0.08 | 5.2±0.01 | 0.5±0.03 | 0.7±0 | 0.8±0.1 | 3.5±0.1 | 4.1±0 |
| *Pellet-AXOS* | 7.1±0.05 | 5.4±0.08 | 5.5±0.06 | 0.2±0.02 | 0.3±0 | 0.3±0.1 | 4.0±0.2 | 4.2±0.3 |
| *WAP-AXOS* | 7.1±0.06 | 6.0±0.06 | 6.0±0.02 | 0.4±0.01 | 0.3±0.05 | 0.4±0 | 3.9±0.3 | 3.7±0.3 |

### Polysaccharide utilizing loci (PUL) in Prevotella copri DSM18205 reveals two putative xylan degrading PULs

To connect the ability to utilize xylan to the enzyme systems in *P. copri*, genes encoding potential xylan degrading enzymes were investigated. The genome of *P. copri* is available (Accession PRJNA30025). According to the polysaccharide utilization-loci database (Terrapon et al. 2018), this microorganism is predicted to encode 17 polysaccharide utilizing loci (PULs). The sizes of the predicted loci range from 20 potential genes (PUL14) to only two-gene PULs (PUL9 and PUL11, including only SusC and SusD homologues, which encode outer membrane proteins that bind and import oligosaccharides, respectively (Koropatkin et al. 2008).

Endoxylanases (EC 3.2.1.8) have according to Cazy (www.cazy.org) been reported in 16 different GH families (3, 5, 8, 9, 10, 11, 12, 16, 26, 30, 43, 44, 51, 62, 98, 141). Analysis of *P. copri* genes shows presence of potential GH encoding genes from these families in six of the 17 predicted PULs (PULs 1, 2, 8, 10, 11 and 15, **Supplementary information, S1**). In PUL1, a gene encoding GH5_4 is present, but this subfamily is reported to encode cellulases (EC 3.2.1.4) and is not likely involved in xylan degradation. PUL2 encodes a potential GH30_3, from which characterized candidates are involved in degradation of β-1,6-glucan linkages (EC 3.2.1.75). PUL8 encodes two putative GH5: GH5_4 (as in PUL1 above) and GH5_7, for which characterized enzymes display endo-β-mannanase activity (EC3.2.1.78). In PUL11, two GH43 subfamilies (4 and 5) are both reported to encode endo-1,5-arabinanases, and this PUL is also not likely involved in degradation of xylan.

This leaves the predicted PUL 10 and 15 as potential xylan degrading loci (**Figure 1**). Both these loci encode enzymes that are putative endoxylanases. Families GH10 and GH5_21 (homologous to the sequences in PUL10) are both reported to encode enzymes with EC3.2.1.8 activity and homologues to the putative GH43_1 and GH10 in PUL15, are reported to encode enzymes with xylosidase (EC3.2.1.37 and endoxylanase (EC3.2.1.8) activity, respectively (www.cazy.org). The genes encoding potential xylan main chain degrading enzymes are in both PULs accompanied by genes that encode enzymes potentially acting on the substituents of various xylan polymers, including potential arabinofuranosidase (PUL10), glucuronidase and galactosidase (both in PUL15). In addition, both PULs encode SusC and SusD homologues, and a gene encoding an inner membrane associated sensor-regulator system, represented by the hybrid two-component systems (HTCS) (Bolam and Koropatkin 2012). In PUL15 a major facilitator superfamily (MFS) transporter gene (Yan 2013) is also present.

**Figure 1.**
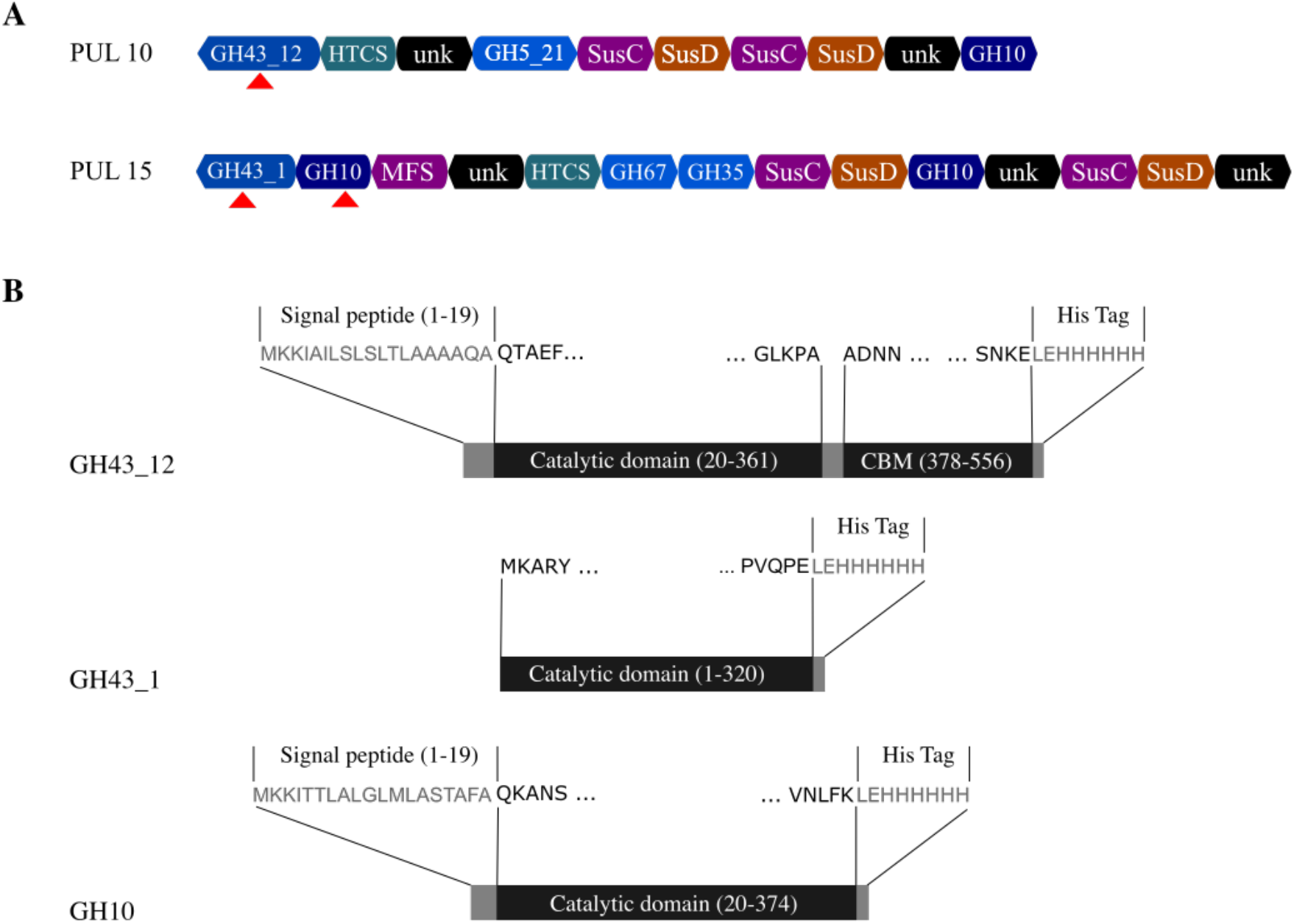
PULs and enzymes characterized. (A) PULs from *P. copri* DSM18205 containing genes potentially involved in xylan degradation (based on *www.cazy.org*). Genes encoding putative GH10 and GH43 enzymes were selected for cloning and expression. (B) The enzymes GH43_12 (PUL 10), GH43_1 (PUL 15) and GH10 (1) (PUL 10) characterized in this work.

### Sequence analysis indicates degradation of arabinoxylan and glucuronoxylan in PUL 10 and 15 of P. copri, using mainly extracellular GHs

All the genes encoding putative GH-enzymes in PUL 10 and 15 are relevant for xylan degradation. PUL 10 contains three genes (with sequence similarities to GH43, GH5 and GH10), and PUL15 contains five genes (with sequence similarities to GH43, GH10 (2 genes), GH67 and GH35), which based on sequence similarities to known GH enzymes encodes xylanases and accessory enzymes (arabinofuranosidase in PUL10 and glucuronidase and galactosidase in PUL15) (**Figure 1, Table 3**). The deduced amino acid sequence of the second putative GH10 in PUL15, termed GH10 (2), was however only 130 amino acid residues corresponding to a molecular mass of 15 kDa, indicating that the gene was fragmented, encoding an incomplete catalytic module. In addition, no signal peptide was predicted for this protein. The deduced amino acid sequences encoded by the remaining four genes were analysed for the presence of signal peptides, showing that all the genes, except the putative β-xylosidase GH43_1 from PUL15, encoded likely signal peptides (**Table 4**). This is suggesting that GH43_1 from PUL15 is located intracellularly in *P. copri*. The presence of signal peptides in the accessory enzymes, also suggest that they may remove substituents extracellularly to facilitate hydrolysis of the polymer and uptake of non-substituted oligosaccharides for intracellular metabolism, which is in accordance with the slower degradation of arabinose, compared to xylose in the growth trials (**Table 2**).

**Table 3.** Annotation and BLAST analysis of the sequences encoded putative enzymes in the polysaccharides utilizing loci (PULs) specialized in xylan degradation in *Prevotella copri* DSM18205.

| PUL | CAZy domain | Putative function | DATABASE UniProtKB<br>Swiss-Prot / TrEMBL |  |  |  | DATABASE UniProtKB<br>Swiss-Prot / PDB |  |  |  | Reference |
| --- | --- | --- | --- | --- | --- | --- | --- | --- | --- | --- | --- |
|  |  |  | Closest Hit | Accession code | Identity (%) | Cover (%) | Closest Hit | Accession code | Identity (%) | Cover (%) |  |
| 10 | **GH10** | β-1,4-xylanase | GH10 of *Prevotella sp.* | A0A3C0D5S0_9BACT | 74.1 | 100 | GH 10 of *Bacteroides intestinalis* DSM 17393 | B3CET4_9BACE | 44.8 | 98.23 | (Zhang et al. 2014) |
| | **GH5_21** | β-1,4-xylanase | GH5 of human gut metagenome | K1RV20_9ZZZZ | 93.8 | 54.76 | Endo-1,4-β glucanase of *Clostridium cellulovorans* | P94622_CLOCL | 29.9 | 27.70 | (Sheweita et al. 1996) |
| | **GH43_12** | α-L-arabinofuranosidase | GH43 of *Prevotella sp.* | A0A354L6I2_9BACT | 99.6 | 100.00 | Xylosidase/arabinosidase of *Butyrivibrio fibrisolvens* | XYLB_BUTFI | 38.2 | 91.19 | (Utt et al. 1991) |
| 15 | **GH43_1** | β-xylosidase | α-N-arabinofuranosidase of *Prevotella copri* | A0A414YDJ3_9BACT | 99.7 | 100.00 | β-xylosidase of *Prevotella ruminicola* B<sub>14</sub> | XYNB_PRERU | 79.9 | 99.38 | (Gasparic et al. 1995) |
| | **GH10 (1)** | β-1,4-xylanase | β-xylanase of *Prevotella copri* | A0A3R5ZWE3_9BACT | 99.7 | 100.00 | Endo-1,4-β-xylanase A of *Prevotella ruminicola* B<sub>14</sub> | XYNA_PRERU | 61.5 | 100.00 | (Gasparic et al. 1995) |
| | **GH67** | α-glucuronidase | α-glucuronidase of *Prevotella sp.* | A0A350PR27_9BACT | 99.1 | 100.00 | Xylan α-(1->2)-glucuronosidase of *Geobacillus stearothermophilus* | AGUA_GEOSE | 43.1 | 85.18 | (Zaide et al. 2001) |
| | **GH35** | β-galactosidase | Uncharacterized protein of *Prevotella copri* | A0A412J314_9BACT | 98.9 | 100.00 | β-galactosidase of *Caulobacter vibrioides* NA1000 | A0A0H3C5N2_C AUVN | 36.00 | 82.30 | (Marks et al. 2010) |
| | **GH10 (2)** | β-1,4-xylanase fragment | GH10 domain of *Prevotella copri* | A0A3R6IT62_9BACT | 97.8 | 70.77 | Endo-1,4-β-xylanase C of *Neosartorya fumigata* | XYNC_ASPFC | 44.4 | 27.69 | (Fedorova et al. 2008) |

**Table 4.** GHs related to xylan hydrolysis encoded within clusters 10 and 15 from *Prevotella copri* DSM 18205 genome according to the CAZy prediction tool. Proteins expressed in soluble form or in inclusion bodies, as well as the presence or absence of signal peptide and molecular weights are shown.

| Cluster | GH Family | Length (aa) | Signal Peptide (SP) | Theoretical MW (kDa) with SP | Theoretical MW (kDa) without SP | Native MW (kDa) | Expression in *E. coli* |
| --- | --- | --- | --- | --- | --- | --- | --- |
| PUL10 | GH10 | 735 | YES (1-22 aa) | 82.4 | 80.1 | - | ND |
|  | GH5_21 | 473 | YES (1-30 aa) | 53.7 | 50.4 | - | Inclusion bodies |
|  | GH43_12 | 556 | YES (1-19 aa) | 61.9 | 60.1 | 77.1 | Soluble * |
| PUL15 | GH43_1 | 320 | NO | - | 36.8 | 77.2 | Soluble ** |
|  | GH10 (1) | 374 | YES (1-19 aa) | 42.7 | 40.7 | 80.1 | Soluble ** |
|  | GH67 | 668 | YES (1-19 aa) | 76.2 | 74.1 | - | Inclusion bodies |
|  | GH35 | 548 | YES (1-19 aa) | 61.7 | 59.6 | - | Inclusion bodies |
|  | GH10 (2) | 130 | NO | - | 14.9 | - | Inclusion bodies |

ND not detected.
\* Expression in *E. coli* BL21(DE3), 25°C, 0.7 mM IPTG, 24 h.
\*\* Expression in *E. coli* BL21(DE3), 37°C, 1.0 mM IPTG, 4 h.

Of the three deduced amino acid sequences of potential xylan degrading enzymes in PUL10, the GH10 candidate is homologous to one of the most well-known endoxylanase families, with more than 3000 candidates reported in the Cazy database (www.cazy.org). PUL10 also encodes a putative xylanase, homologous to GH5_21. GH5_21 is reported as a subfamily of GH5 that encode enzymes with EC 3.2.1.8 activity (Aspeborg et al. 2012). To date (june 2020), 35 sequences of GH5 subfamily 21 are present in the CAZy database and all of them belong to the Bacteroidetes. From those, only four have been biochemically characterized, including two *Prevotella bryantii* B_1_4 enzymes, which showed activity against wheat arabinoxylan (Dodd et al. 2010). The deduced amino acid sequence of the *P. copri* GH5_21 from PUL 10 showed > 92 % of identity with GH5 detected in human gut metagenome and other *P. copri* strains, according to a BLAST analysis against UniProtKB Swiss-Prot/TrEMBL database (**Table 3**). PUL 10 also contains a sequence of a putative GH43_12 enzyme, which showed 38.2 % of identity with a characterized enzyme of *Butyrivibrio fibrisolvens,* according to a BLAST analysis against UniProtKB Swiss-Prot/PDB database (**Table 3**). Accordingly, the *B. fibrisolvens* GH43 from subfamily 12 was reported to present α-L-arabinofuranosidase (EC 3.2.1.55) activity (Mewis et al. 2016).

The four complete genes in PUL15 displayed sequence similarities to GH43, GH10, GH67 and GH35. The potentially intracellularly located GH43 candidate in this PUL is similar to subfamily 1. Homologues to GH43_1 and GH10 in PUL15 were reported to encode enzymes with β-xylosidase (EC 3.2.1.37) and endo-β-xylanase (EC 3.2.1.8) activity, respectively. For example, GH43_1 showed 79.9 % sequence identity with a β-xylosidase of *Prevotella ruminicola* B_1_4 (**Table 1**).Sequences homologous to the GH67 in PUL 15, has been reported to encode enzymes with α-glucuronidase (EC 3.2.1.139) activity (Malgas et al. 2019). The translated protein sequence showed 43.1% of identity with a xylan α-(1,2)-glucuronosidase from *Geobacillus stearothermophilus* (**Table 3**). PUL15 also contains a potential β-galactosidase, with sequence similarities to GH35. Interestingly, many types of xylans from agricultural resources are reported to contain minor amounts of galactose (*e.g*. sugar cane bagasse, L Khaleghipour, J.A. Linares-Pastén^a^, H Rashedi, S.O.R. Siadat, A. Jasilionis, S. Al-Hamimi, R.R.R. Sardari, E. Nordberg Karlsson, manuscript), making this activity relevant in a xylan degradation PUL.

### Cloning and production of the xylan degrading enzymes from P. copri

Synthetic sequences encoding the GHs were cloned into pET-21b (+) plasmid and transferred to *E. coli*. This resulted in the production of soluble active forms from three out of the seven putative enzymes cloned: GH43_12 from PUL 10 and GH10 (1) and GH43_1 from PUL 15. The theoretical molecular weights of the two PUL 15 enzymes were of 37 kDa (GH43_1) and 41 kDa (GH10) (**Table 4**), which was in agreement with their apparent molecular weights estimated by SDS-PAGE (**Supplementary Information S2**). Based on the size of the catalytic modules of the respective families, both are single module enzymes comprising one catalytic module each. The native molecular mass analysed by SEC showed that both enzymes were present in solution as dimers. On the other hand, the β-xylosidase GH43_12 from PUL 10 was confirmed to be a monomer in solution, with a molecular mass of 60.1 kDa (**Table 4**) that denoted a two-domain enzyme (also see modelling results below).

Most commonly, GH43 enzymes that show β-xylosidase activity have been reported to have dimeric (or, in some cases tetrameric) structures; such as RS223-BX enzyme from an anaerobic mixed microbial culture. This enzyme showed an activation with the addition of divalent cations and their removal caused changes in the quaternary structure of the enzyme (Lee et al. 2013). Another dimeric GH43 β-xylosidase from *Rhizophlyctis rosea* (RrXyl43) exhibited a dimeric structure, presenting Ca^2+^ and Na^+^ ions in its proposed model that further supports such structure. However, the enzyme remained partially active as a monomer (Huang et al. 2019).

### Biochemical characterization of the recombinant proteins

The enzymes were screened for activity using aryl substrates and xylans of different origins. Both GH43_12 from PUL10 (potentially extracellular) and GH43_1 from PUL 15 (potentially intracellular) showed exo-acting activity resulting in hydrolysis of *p*NP-arabinofuranoside and *p*NP-xylopyranoside substrates.

In accordance with previously data for subfamily 12 (Kobayashi et al. 2020; Mewis et al. 2016), the specific activity for GH43_12 was most significant on *p*NP-Ara (1.7 U. mg^−1^), value comparable to its specific activity on arabinoxylan (1.2 U. mg^−1^) (**Table 5**). GH43_12 showed little activity on arabinans, suggesting a low preference on arabinoside 1,5-linkages, dominant in this substrate. However, the enzyme produced a complete conversion of 1,2-(A^2^XX) and 1,3-(XA^3^XX) arabino substituted AXOs after 30 minutes of reaction, but no activity was detected on 1,2- / 1,3- double substituted (A^2,3^XX) AXOs (**Figure 2**). Thus, GH43_12 is concluded to display a debranching function on AXOS with single substituted arabinose groups.

**Figure 2.**
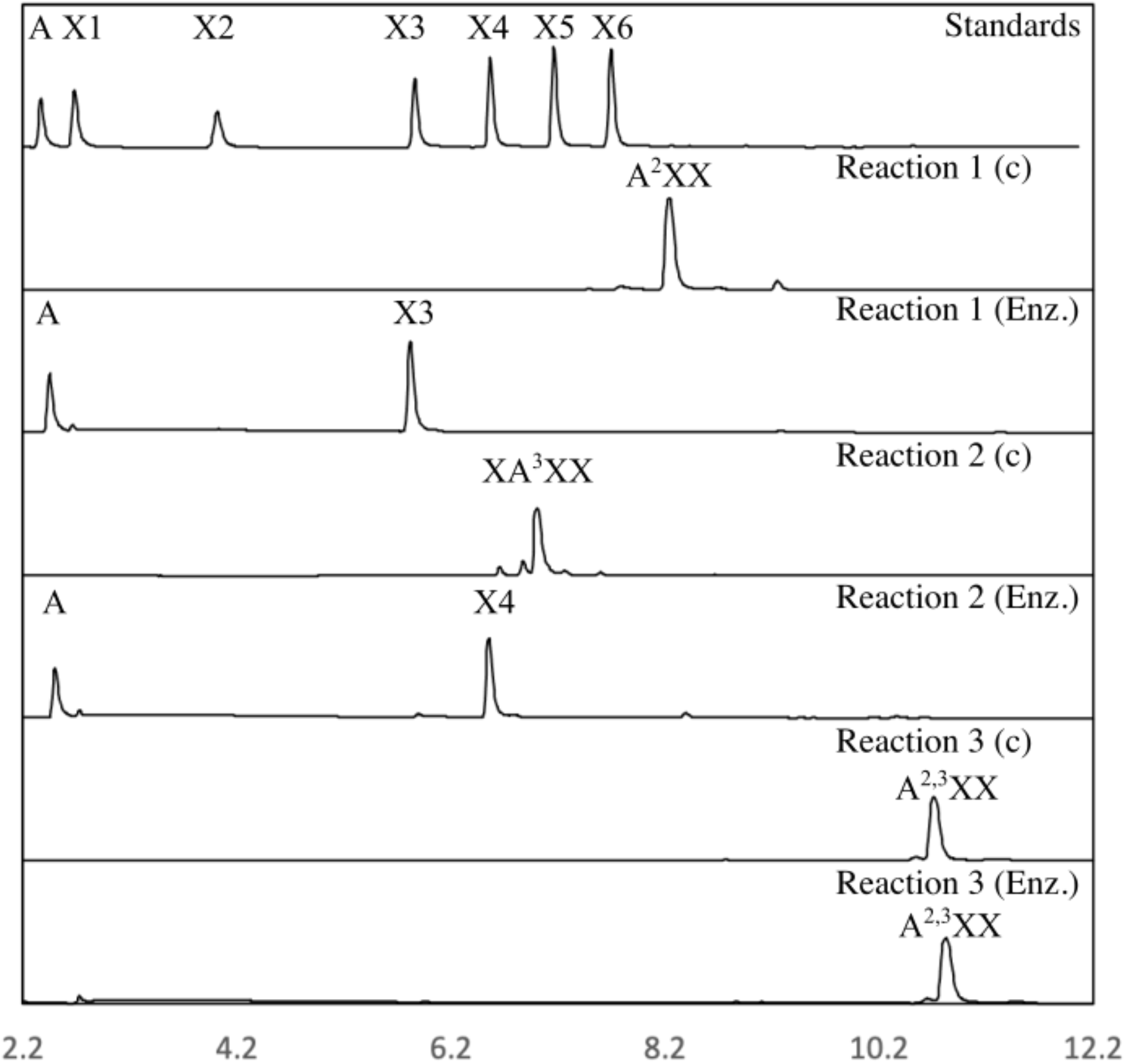
Product profiles of GH43_12 reaction against single and double arabino substituted AXOS. The standards are arabinose (A), xylose (X1), xylobiose (X2), xylotriose (X3), xylotetarose (X4), xylopentaose (X5) and xyloexaose (X6). The substrates used were single arabino substituted AXOs (A^2^XX, and XA^3^XX) and double AXOs (A^2,3^XX). The enzyme showed activity on the single substituted substrates, but no activity is detected on the double substituted one. Enz.= substrate after incubation with GH43_12, c= substrate control

**Table 5.** Screening of specific activities of the three produced xylan-acting enzymes on both, synthetic substrates (*p*-NP-glycosides) and different xylans.

| Substrate | *GH43_12 (PUL 10)*<br>(U. mg<sup>-1</sup>) | *GH10 (1) (PUL 15)*<br>(U. mg<sup>-1</sup>) | *GH43_1 (PUL 15)*<br>(U. mg<sup>-1</sup>) |
| --- | --- | --- | --- |
| *p*-NP-β-D-Xyl1 | 0.1 ± 0.0 | ND | 1.3 ± 0.1 |
| *p*-NP-β-Xyl2 | 0.6 ± 0.1 | 207.8 ± 43.2 | 1.7 ± 0.1 |
| *p*-NP-β-Xyl3 | 0.5 ± 0.0 | 300.7 ± 42.2 | 1.4 ± 0.2 |
| *p*-NP-α-L-Ara | 1.7 ± 0.3 | ND | 1.0 ± 0.1 |
| *p*-NP-β-D-Glu | ND | ND | ND |
| *p*-NP-β-D-Gal | ND | ND | ND |
| *p*-NP-β-D-Man | ND | ND | ND |
| Birchwood Xylan | 0.8 ± 0.0 | 290.8 ± 10.1 | 13.7 ± 0.4 |
| Beechwood Xylan | 0.8 ± 0.2 | 611.7 ± 28.1 | 23.3 ± 3.3 |
| Arabinoxylan | 1.2 ± 0.1 | 258.5 ± 12.2 | 10.9 ± 1.5 |
| Quinoa stalks Xylan | 0.5 ± 0.0 | 449 ± 30 | 15.7 ± 0.5 |
| Arabinan Sugar Beet | 0.3 ± 0.0 | ND | ND |
| Debranched Arabinan | 0.4 ± 0.0 | ND | ND |

ND not detected.

The specific activity of the intracellularly located GH43_1 (PUL 15) on *p*-NP-Ara and *p*-NP-Xyl was alike, coinciding with a dual specificity previously reported for some enzymes in this subfamily (Huang et al. 2019; Matsuzawa et al. 2017; Mewis et al. 2016). This enzyme presented a specific activity over xylan substrates higher (~ one order of magnitude) than those of GH43_12 (**Table 5**). As this enzyme hydrolysed xylan and xylooligosaccharides to xylose, we considered that, at least *in vitro*, is not limited to substrates of short degree of polymerization (DP). Yet, its potential intracellular location makes it likely that the native function of the evaluated GH43_1 is the degradation of xylooligosaccharides to xylose in *P. copri*. The GH10 (1) enzyme (PUL 15) displayed the highest specific activity against various xylan substrates assayed but was also active on the aryl-substrates with a minimum DP of 3 (*p*-NP-xylobiose). No activity was observed on *p*NP-xyloside, indicating that this GH10 (1) lacked exo-activity. No activity on *p*-NP-glucoside, *p*-NP-galactoside or *p*-NP-mannoside was detected for any the enzymes (**Table 5**).

Further studies of temperature and pH optima were made utilizing as substrates those showing the highest specific activity for each enzyme. Hence, *p*-NP-arabinofuranoside was chosen for GH43_12 from PUL 10, while beechwood xylan was used as a model substrate for both enzymes from PUL 15. Despite analysing two putatively extracellular enzymes (GH43_12; PUL 10 and GH10; PUL 15) and one intracellular enzyme (GH43_1; PUL 15), no major differences of temperature profiles were observed (**Figure 3**).

**Figure 3.**
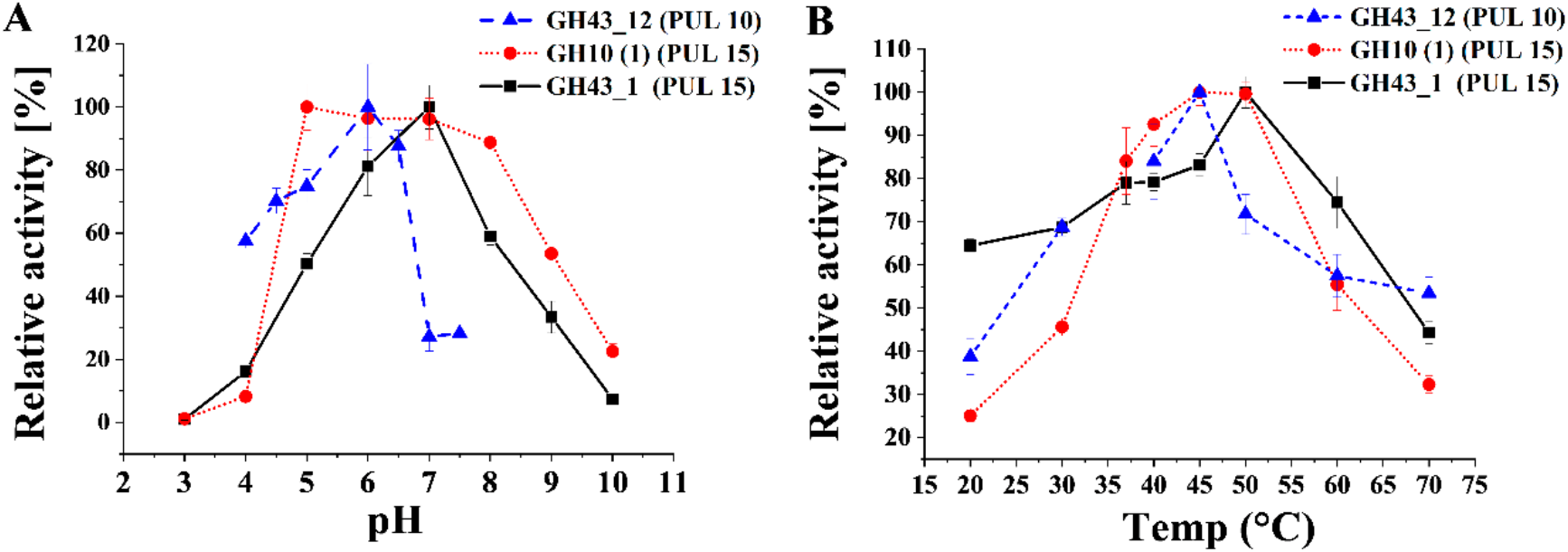
Relative activity at different pH (**A**) and temperatures (**B**) of the cloned enzymes. GH43_12 (PUL 10) was assayed with *p*NP-arabinofuranoside as substrate. GH10 (1) and GH43_1 (PUL 15) were evaluated with beechwood xylan.

It has been hypothesized that extracellular enzymes are active in a wider range of conditions (pH and temperature) than intracellular ones (Chang et al. 2017; Mechelke et al. 2017), but no differences were detected here outside the slightly lower apparent optimum temperature of GH43_12; PUL10 which was analysed using an aryl substrate. Instead, the pH profile was narrower (with a lower pH optimum) than expected. As an overall observation, the pH and temperature profiles may indicate that the enzymes assayed are well suited for activity in the gut, acting at neutral pH and with high activity at 37°C.

### Hydrolysis product profiles

Hydrolysis products were analysed using four different xylan substrates (birchwood xylan, beechwood xylan, arabinoxylan, and quinoa stalks xylan (**Figure 4**). For this purpose, focus was put on the PUL 15 enzymes, as GH43_12 (from PUL 10), had its highest specific activity on the short aryl-substrate *p*-NP-Ara, resulting in release of monosaccharides.

**Figure 4.**
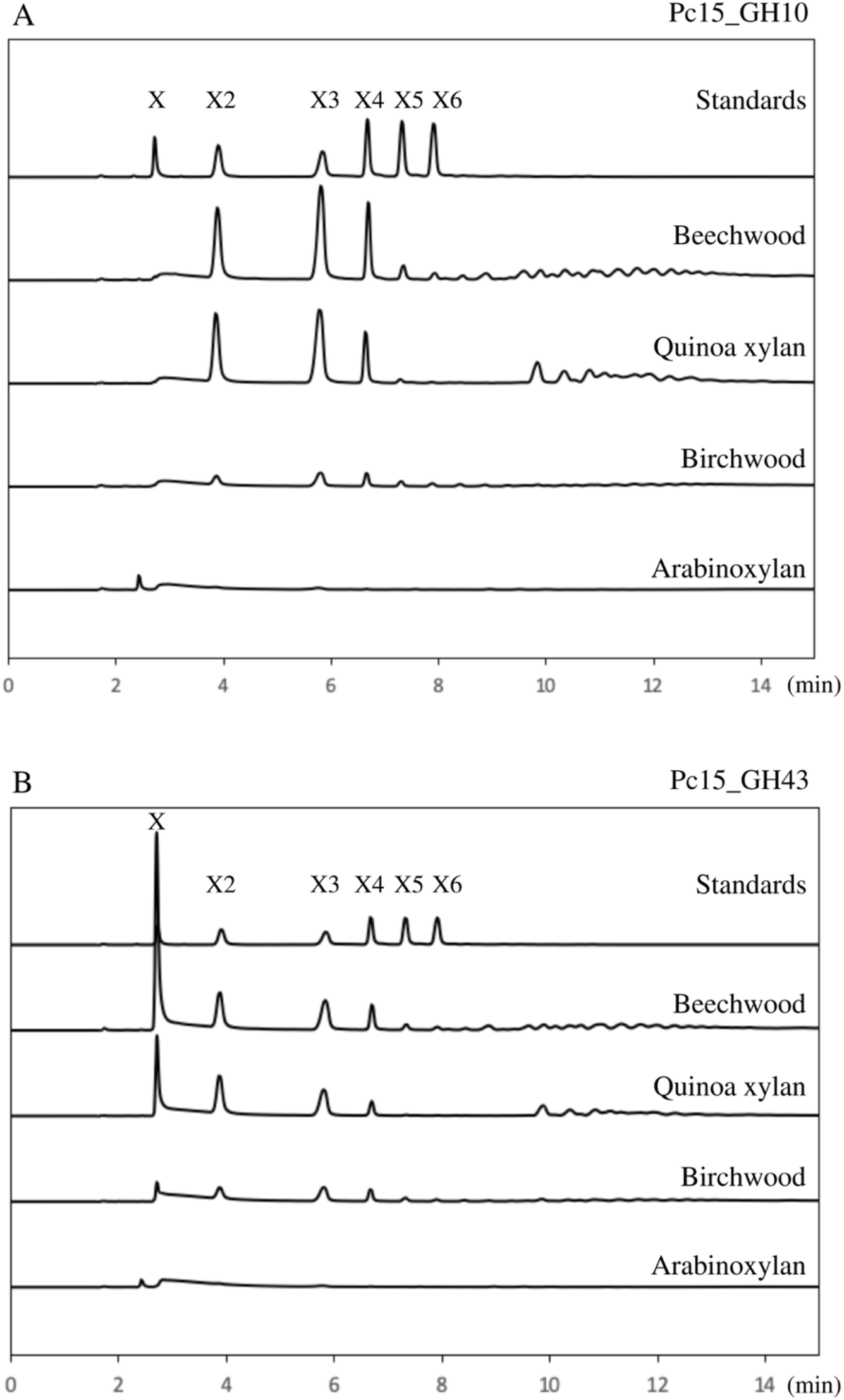
Product profiles of *P. copri* DSM18205 enzymes on different type of xylan, anlyzed by HPAEC-PAD. (A) GH10 (1) fromPUL 15 (abbreviated Pc15_GH10) and (B) GH43_1 from PUL 15 (abbreviated Pc15_GH43). The standards are xylose (X1), xylobiose (X2), xylotriose (X3), xylotetraose (X4), xylopentaose (X5) and xylohexaose (X6). Despite the profiles are different, both enzymes showed highest activity on beechwood, followed by quinoa xylan, birchwood and arabinoxylan (see also **Table 3**).

For all xylan substrates, hydrolysis products were obtained using both GH10 and GH43_1 from PUL 15 (**Figure 4**) using the non-hydrolysed substrates as control. When birchwood, beechwood or quinoa xylans were utilized, GH10 (1) produced X2, X3 and X4 as major products, whereas xylose was the main product from GH43_1 (**Figure 4**). This confirms the endoxylanase activity of GH10 from PUL 15. Low amounts of hydrolysed products were observed with arabinoxylan as substrate, exposing the need of additional debranching enzymes for an effective degradation of this substrate. Release of xylose by the action of the GH10 enzymes has previously been reported for related endo-xylanases, evidencing ability to act on low molecular weight XOS (Chapla, Pandit, and Shah 2012; Rahmani et al. 2019; Salas-Veizaga et al. 2017). The apparent lack of generation of xylose by GH10 (1), despite presence of xylooligosaccharides with a DP between 2 to 5 (**Figure 4A**), gives to the GH10 (PUL 15) endo-1,4-β-xylanase from *P. copri* DSM18205 a significant biotechnological potential for oligosaccharides production with prebiotic activity (Linares-Pasten, Aronsson, and Karlsson 2018; Rahmani et al. 2019; Samanta et al. 2015).

The main product of GH43_1 from PUL 15 was instead xylose, but in addition to xylose significant amounts of oligosaccharides of DP2, 3 and 4 were also observed for this enzyme on all xylan assayed, despite its potential intracellular location. Noticeable, there was only limited hydrolysis observed for the arabinoxylan substrate (from rye flour), corresponding to a diffuse wide peak observed in the HPAEC chromatograms (**Figure 4B**) indicating that this substrate could not be hydrolysed by the enzyme. This indicated that arabinoxylan is hardly hydrolysed by this enzyme.

The profile of xylan hydrolysis products by GH43_1 (PUL 15) (**Figure 4**) is in accordance with exo-β-xylanase activity, which is also supported by its activity on aryl substrates, indicating β-xylosidase and α-L-arabinofuranosidase activities observed in the substrate specificity studies (**Table 5**). Typical β-xylosidases act on low DP xylooligosaccharides to produce xylose as the last product; however, GH43_1 also acts on polymeric substrates, yielding significant amounts of xylose as final hydrolysis product thus suggesting an exo-β-xylanase mode of action. A similar mechanism of action was reported by a multiple activity GH43 enzyme (exo-β-xylosidase, endo-xylanase, and α-L-arabinofuranosidase) from *Paenibacillus curdlanolyticus* B-6, GH43B6, which initially produced X5 and X4 from X6 and then, xylose as the final hydrolysis product (Wongratpanya et al. 2015).

### Enzyme kinetic studies

Measurement of kinetic parameters for the two PUL15 candidates were based on the initial rate of product formation from xylooligosaccharides and aryl substrates. The resulting data were suitable for a nonlinear regression analysis according to the Michaelis–Menten equation (**Supplementary Figure 3**).

Kinetic parameters of the purified GH43_12 from PUL 10 using *p*-NP-Ara as a substrate resulted in a ***Km*** of 3.2 ± 0.3 mM, ***kcat*** of 7.5 s^−1^, and a ***kcat/Km*** of 2.4 s^−1^ mM^−1^ (**Table 6**). Among GH43 enzymes with α-L-arabinofuranosidase activity diverse kinetic parameters were reported for the *p*-NP-Ara hydrolysis. In the case of GH43_12 from *P. copri* DSM18205, kinetic studies revealed that the ***Km*** value was 7.2- and 1.2-fold lower (assuming higher affinity) than those previously described for enzymes AxB8 from *Clostridium thermocellum* B (de Camargo et al. 2018), and Xsa43e from *Butyrivibrio proteoclasticus* B316 (Till et al. 2014), respectively. The ***kcat/Km*** value was 1.87-fold lower than the one determined for the AxB8 enzyme, but 12-fold higher than that observed for Xsa43e. This *B. proteoclasticus* enzyme, Xsa43e, mainly showed activity on low DP AXOS, contributing to arabinoxylan debranching, as was also observed for the *P. copri* GH43_12.

**Table 6.** Kinetic constants of GH43_12 (PUL 10), GH10 (1) (PUL 15) and GH43_1 (PUL 15) determined at 37 °C and pH 5.5.

| Enzyme | Substrates | Kinetic constants |  |  |  |
| --- | --- | --- | --- | --- | --- |
|  |  | *V<sub>max</sub>*<br>(*U. mg*<sup>-1</sup>) | *K<sub>m</sub>*<br>(*mM*) | *k<sub>cat</sub>*<br>(*s*<sup>-1</sup>) | *k<sub>cat</sub>/K<sub>m</sub>*<br>(*s*<sup>-1</sup> *mM*<sup>-1</sup>) |
| GH43_12<br>(PUL 10) | p-NP-α-L-arabinofuranoside | 7.3 ± 0.2 | 3.2 ± 0.3 | 7.5 | 2.4 |
| GH10 (1)<br>(PUL 15) | p-NP-β-xylobioside | 371 ± 13 | 1.0 ± 0.1 | 264.0 | 275.0 |
| | p-NP-β-xylotrioside | 1128 ± 80 | 2.6 ± 0.6 | 802.8 | 302.9 |
| GH43_1<br>(PUL 15) | 1,4-β-D-Xylobiose | 39.8 ± 2.3 | 9.2 ± 1.6 | 24.4 | 2.7 |
| | 1,4-β-D-Xylotriose | 21.2 ± 1.2 | 5.5 ± 1.1 | 13.0 | 2.4 |
| | 1,4-β-D-Xylotetraose | 3.5 ± 0.2 | 0.6 ± 0.2 | 2.1 | 3.7 |
| | 1,4-β-D-Xylopentaose | 5.4 ± 0.2 | 0.5 ± 0.1 | 3.3 | 6.3 |
| | 1,4-β-D-Xylohexaose | 1.9 ± 0.2 | 0.8 ± 0.4 | 1.1 | 1.4 |

The ***Km*** value of GH10 (1) (PUL 15) was 1.0 mM for p-NP-Xyl2 and 2.6 mM for *p*-NP-Xyl3, demonstrating a better affinity for short-chain aryl substrates (**Table 6**). But, the turnover number (***kcat***) and the catalytic efficiency ***kcat/Km*** for *p*-NP-Xyl3 were 3-fold and 1.1-fold higher than those for *p*-NP-Xyl2, respectively. Overall, this indicates that longer-chain aryl substrates are preferred by this enzyme, in line with an endo-acting activity.

The enzyme GH43_1 (PUL 15) it showed higher ***Km*** values (lower affinity) for X2 and X3 compared to those for X4 – X6 (**Table 6**). This may denote the presence of at least 4 distinct subsites in the enzyme. The turnover was the highest for X2 (followed by X3), result that combined with the preference for producing xylose indicates that the −1, +1 and +2 subsites could be accountable for high affinity binding. The catalytic efficiency (***kcat/Km***) was however highest for the X5 oligosaccharide (6.3 ***s^−1^mM^−1^***) which may suggest a faster release of the product from the enzyme. The presence of multiple subsites could explain the possible exo-β-xylanase of this enzyme on polymeric substrates (**Table 5**), making GH43_1 activity greatly divergent from other β-xylosidases present in microorganisms with prebiotic characteristics isolated from the GI tract (Lasrado and Gudipati 2013).

### Predicted molecular structures

The three-dimensional structures of characterized enzymes, GH43_12; GH43_1 and GH10 (1) were obtained by homology modelling using the combination of several crystallographic templates. The overall quality of the obtained models varied from satisfactory to good, indicating that the models are reliable (**Supplementary Information S4**),.

GH43_12 (PUL10) is an enzyme with relatively low specific activity. The enzyme, however, showed debranching activity on AXOS as well as a higher activity on *p*NP-a-arabinofuranoside and arabinoxylan than that on *p*NP-xylosides and xylans with low arabinosylation (**Table 5**). This clearly indicates that GH43_12 is an arabinofuranosidase, in accordance with what was proposed for other characterized members of this subfamily, with a role in debranching arabinoxylan or arabinoxylooligosaccharides (**Table 3**). The presence of an N-terminal signal peptide (Met1 to Ala19), implying that the enzyme follows a secretion path, which is consistent with activity on large substrates such as arabinoxylan. The protein was studied excluding the signal peptide and was observed to be composed by two-domains according to the molecular model (in line with the sequence analysis), with the catalytic domain at the N-terminal region and a putative carbohydrate binding module (CBM) the in the C-terminal (**Figure 5A**). This was in accordance with the predicted and experimentally observed molecular mass of the enzyme. The catalytic domain has the typical 5-fold β-propeller 3D-structure of the family GH43 while the C-terminal domain has a β-sandwich fold characteristic of the family CBM6.

**Figure 5.**
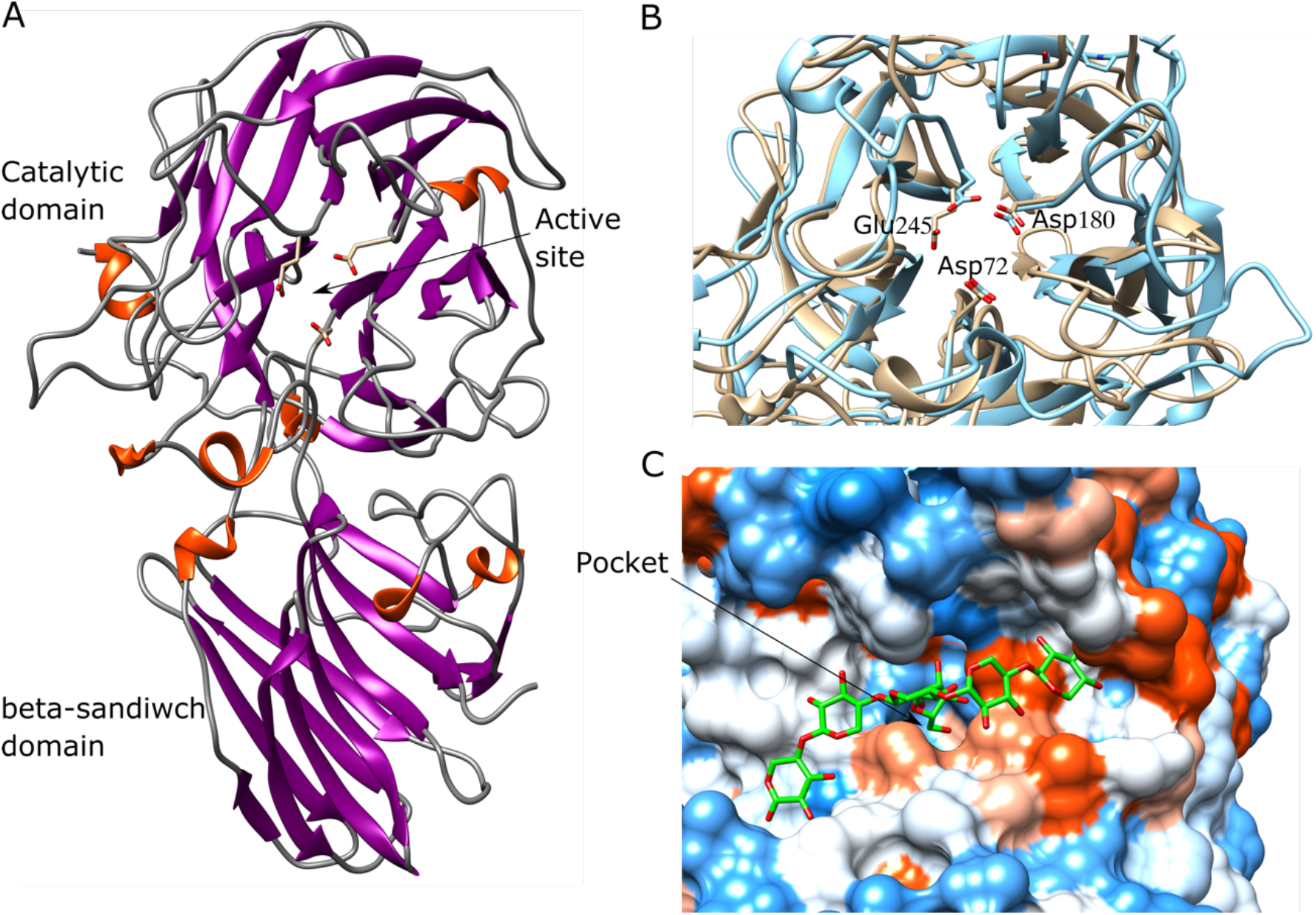
Molecular model of arabinofuranosidase GH43_12 from PUL 10. (A) Overall structure. (B) Conserved catalytic triad. Overlapped model (in brown) and crystallographic structure (in cyan) of *Lactobacillus brevis* arabinofuranosidase (PDB: 5M8B). (C) Active site surface. Grove with a central pocket where the catalytic triad is located surrounding the arabinoside moiety. The ligand docked corresponds to a XXA^2^XX oligosaccharide (represented in green).

The active site is a grove containing a pocket where the catalytic amino acids are located (**Figure 5B**). This suggests that the xylan backbone would bind in the grove with the arabinofuranoside branch into the pocket. The catalytic amino acids are conserved and can be clearly predicted overlapping the model with the crystallographic structure of an arabinofuranosidase GH43, for instance *Lb*Araf43 (PDB: 5M8B) (Linares-Pastén et al. 2017). Thus, Asp57 is predicted as catalytic base, Glu230 as catalytic proton donor and the residue Asp165 as the residue suggested to aid in modulating the pKa of the proton donor (**Figure 5C**) (Nurizzo et al. 2002).

The C-terminal domain has an unknown function. The similarity in fold to CBM6, suggests that this domain may contribute to bind a xylan backbone. However, this remains to be proven. Both experimental as well as computational studies suggest that GH43_12 would be an arabinofuranosidase specialized in debranching arabinose moieties from arabinoxylan polymers in the extracellular medium.

GH43_1 (PUL15) is an enzyme that has shown activity on *p*NP-α-arabinofuranoside and arabinoxylan as well as on *p*NP-xylosides and surprisingly (considering its intracellular location) also on xylan (**Table 3**). The highest activity was, however, observed on xylooligosaccharides, followed by beechwood xylan (which, however, according to our analysis was an oligomeric substrate, see **Supplementary Information S5**) and the lowest on arabinoxylan, indicating preference for non-arabinosylated xylan backbones. The overall 3D structure is a single domain catalytic module with a 5-fold β-propeller fold (**Figure 6A**). The predicted active site is a relatively deep pocket formed with a significant contribution of the loop-helix-loop Asp33 to Gln46 (**Figure 6B**). The shape of the active site would suggest an exoglycoside active enzyme. The kinetic analysis showed a decrease in *K_m_* with increasing length of the substrate (from DP2 to DP4), indicating 4 subsites in the active site, which correlates with the deep pocket shape of the active site. However, deeper studies on mechanistic aspects are required to explain the presence of oligosaccharides as hydrolysis products. The catalytic triad is well conserved and can be predicted by overlapping with crystallographic structures of glycoside hydrolases form the family GH43. Thus, the Asp14 corresponds to the catalytic base, Glu221 to proton donor and the residue Asp135 to the suggested to aid in modulating the pKa of the proton donor (**Figure 6C**).

**Figure 6.**
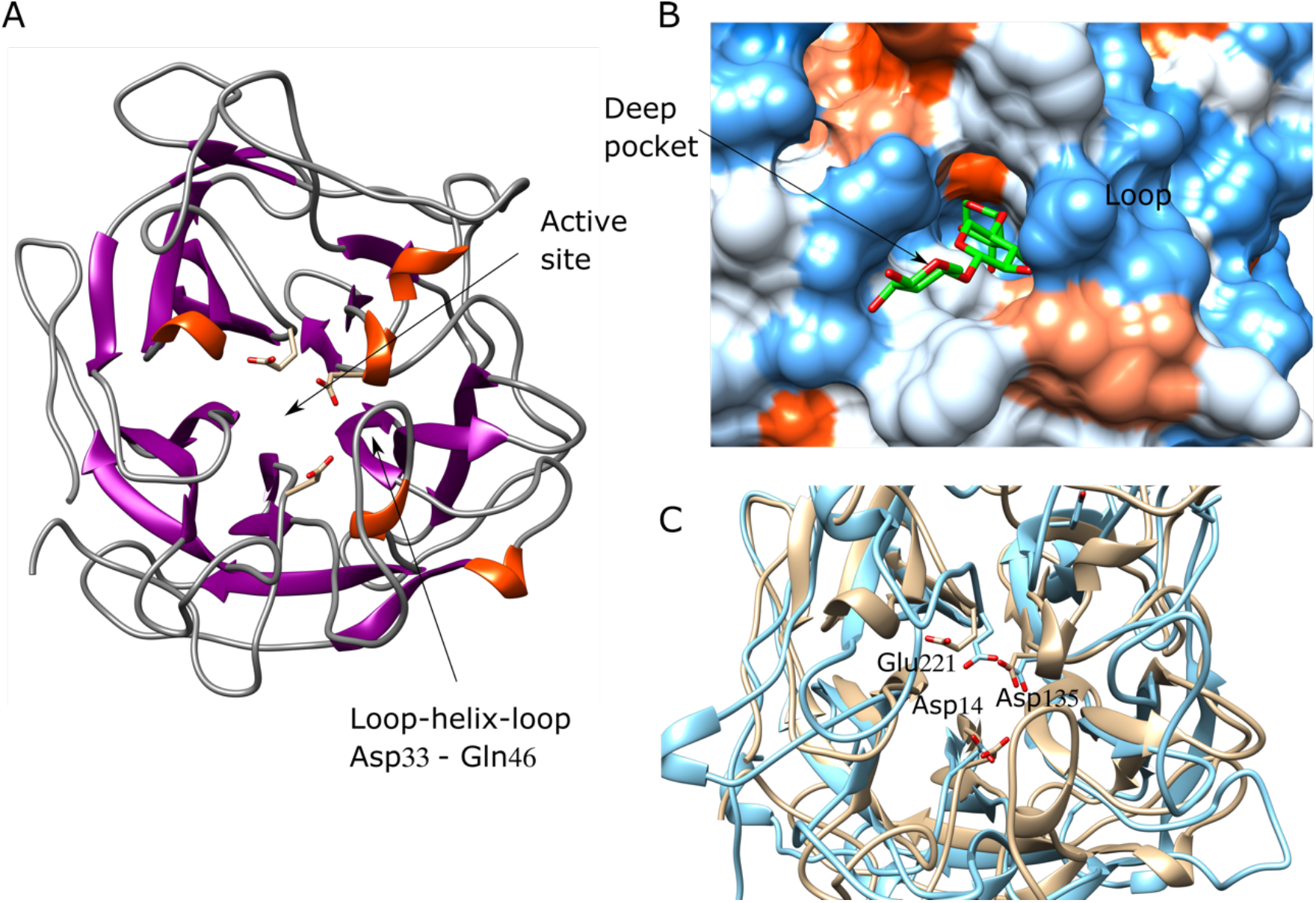
Molecular model of xylosidase/arabinofuranosidase GH43_1 from PUL 15. (A) Overall structure. (B) Active site surface. Deep pocket containing the ligand xylotriaose where the catalytic triad is located. (C) Conserved catalytic triad. Overlapped model (in brown) and crystallographic structure (in cyan) of *Lactobacillus brevis* arabinofuranosidase (PDB: 5M8B).

GH10 (1) (PUL15) has endoxylanase activity with highest activity on the oligomeric beechwood xylan and lowest on pNP-1,4-β-xylobiose (**Table 5**). This enzyme has a signal peptide and secretion is expected. Therefore, its activity on polysaccharides can take place in the extracellular medium. The molecular model is a 3D structure (α/β)_8_-TIM barrel, characteristic of family GH10 (**Figure 7A**). Family GH10 xylanases has a catalytic mechanism with retention of configuration of the anomeric carbon in the substrate. In this enzyme the Glu161 was predicted as the general/acid base, and the Glu266 as catalytic nucleophile (**Figures 7A and 7C**). This model has a good quality z-score (**Supplementary Information S4**), which also allowed reliable prediction of interactions with ligands and acceptance of substituents both in the glycone as well as aglycone subsites (**Figure 7B-C**). Thus, 3 glycone subsites and 4 aglycone subsites were predicted. Some of the xylose subunits of the ligands have hydroxyl groups exposed to the solvent, therefore we can speculate that these groups can be substituted, for instance with arabinofuranose (Araf) or methyl glucuronic (MeGlcA) acid moieties. In the computational model obtained, the hydroxyl groups in subsites −3, +1, +3 and +4 are exposed to the solvent (**Figure 7B**), which indicates the acceptance of substituents (**Table 7**). In general, endoxylanases GH10 can accept substrates more substituted than GH11 (Linares-Pasten, Aronsson, and Karlsson 2018; Nordberg Karlsson et al. 2018). Thus, the different activities on xylans (**Table 4**) depend of the degree and pattern of substitution. These results may explain the low activity of this GH10 in arabinoxylan compared to beechwood xylan, probably due to a high degree of decoration (~ 40% substitution by arabinosyl residues) of the xylan backbone chain when compared to beechwood xylan (~ 13% substitution by glucuronic acid residues). According to the computational analyses, GH10 (1) from PUL 15 requires two unsubstituted xylose residues before the cut-off site for hydrolysis, due to that subsites −2 and −1 cannot accept substituents (Ara*f*/MeGlcA) on the xylose chain (**Table 7**).

**Figure 7.**
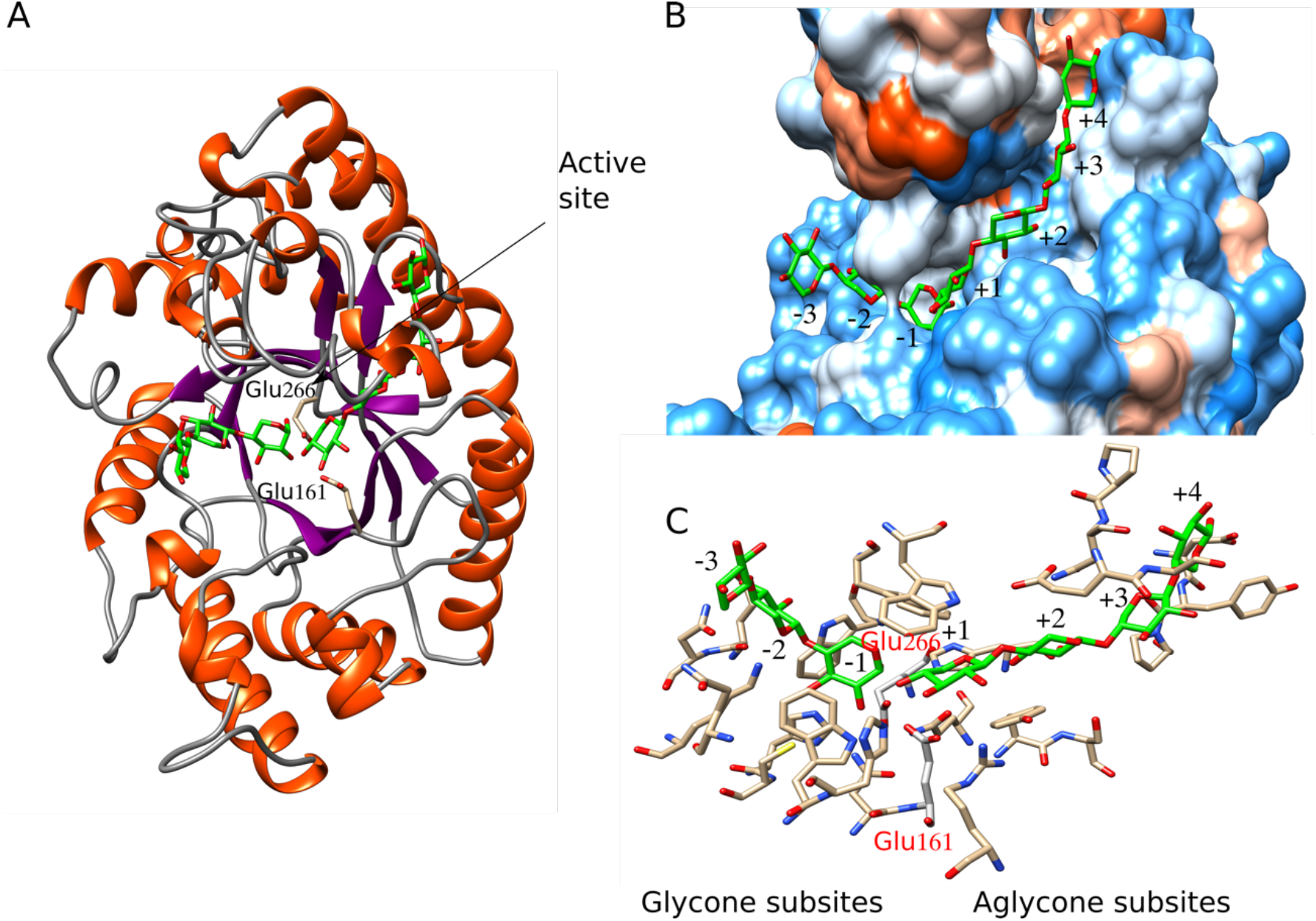
Molecular model of the complex 1,4-β-endoxylanase GH10 (1)-ligands. (A) Overall structure. (B) Active site surface. Seven subsites were predicted, xylose units from the ligand have hydroxyl groups exposed to the solvent in subsites −3, +1, +3 and +4. (C) Detail of the residues surrounding every predicted subsite (see also table 5). Glu266 is predicted as catalytic nucleophile while Glu161 as acid/base residue.

**Table 7.** Molecular model of GH10 (1) from PUL 15, including residues surrounding every predicted subsite and potential substituents allowed in the subsites.

| Substituent | Glycone subsites |  |  | Aglycone subsites |  |  |  |
| --- | --- | --- | --- | --- | --- | --- | --- |
|  | -3 | -2 | -1 | +1 | +2 | +3 | +4 |
| Araf / MeGlcA | P | B | B | P | B | P | P |
| Residues (see also **Figure 7C)** | Gln115 | Glu71,<br>Asn72,<br>Lys175 | His108,<br>Cys109,<br>Trp112,<br>Asn160,<br>Glu266*,<br>Trp332 | Glu161*,<br>Arg170,<br>Tyr204,<br>Gln235,<br>His237,<br>Trp340,<br>Trp344 | Ser205,<br>Glu276 | Pro243,<br>Asn274,<br>Gly277 | Asp241,<br>Try242,<br>Pro273 |

\*Predicted catalytic residues. P=permitted, B=banned,

The amount and conservation of subsites present in this enzyme (**Figure 7C** and **Table 7**) could also explain the activity results observed against the aryl substrates. The enzyme showed strong interactions between the aryl substrates and the −1 and −2 subsites of the glycone region. However, the activities observed against *p*NP-Xyl3 and *p*NP-Xyl2, showed a modest difference, implying that the third binding site in the glycone region (−3) interacts weakly with the substrate. Pell et al. (2004), in their study of a GH10 of *Cellvibrio japonicus*, observed that subsite −2 of catalytic site showed a similar free binding energy (*ΔG*) to the other subsites analyzed: −3 and +2. These authors concluded that the poor activity of GH10 from *C. japonicus* against xylooligosaccharides is the result of a compromised −2 subsite.

## Discussion

The mammalian gut constitutes a highly complex and competitive ecosystem with a strong selective pressure for the microorganisms inhabiting it. In this sense, Bacteroidetes members, the dominant phylum in mammalian gut, were described to have a highly flexible metabolism, being able to alter the gene expression to adapt to the changes of substrate availability in their environment (Haller 2018; Sonnenburg 2005; Yadav et al. 2018). Bacteroidetes genomes generally encode CAZYmes in abundance, especially glycoside hydrolases and polysaccharide lyases (El Kaoutari et al. 2013) that stems from the outer membrane-localized enzymatic complexes or are secreted via a signal peptide (Accetto and Avguštin 2015). Many of such enzymes are aimed to degrade dietary fibres, for which further fermentation is a critical process for the function and integrity of both the bacterial community and the host cells. Moreover, a unique feature of this phylum is that genes encoding various carbohydrate active enzymes, proteins and transporters required for saccharification of complex carbohydrates are organized in PULs, allowing a better utilization of the substrates. It has been observed a degree of the plasticity of the PULs repertoires due to lateral gene transfer, as was described for PULs specialized in porphyran degradation from a marine Bacteroidetes isolated in Japan, probably due to consumption of non-sterile red algae (Hehemann et al. 2010). Further studies on genomic and metagenomic data from mammalian gut and rumen, showed that PUL variants may be specialized for specific representatives of broad substrate classes (Flint et al. 2012; Johnson et al. 2017; Thomas et al. 2011).

The presence of PULs encoding specialized enzymes in the degradation of different types of xylan is a characteristic shared in species of *Prevotella* and *Bacteroides*. It was observed that *P. copri* DSM 18205 displays a larger proportion and diversity of CAZymes (and PULs) than other *Prevotella* species isolated from gut samples. Such metabolic capacity to degrade carbohydrates was shown to be comparable to that of rumen isolated strains (*e.g. P. bryanti*) including strains inhabiting the oral cavity (*P. buccae*) (Accetto and Avguštin 2015).

In the present work, two PULs (10 and 15) from *P. copri* DSM 18205 were detected to encode enzymes aimed at xylan utilization. Three of those enzymes, GH43_12 from PUL 10 and GH10 (1) and GH43_1 from PUL 15, were successfully cloned, over-expressed and characterized. These enzymes present unique and, possibly synergistic, features in the hydrolysis of the xylan chain and its substituent groups.

The analysis of the extracellular GH10 (1) in PUL 15 from *P. copri* showed a broad substrate specificity (**Table 5**), a common feature among endo-β-xylanases in the GH10 family (Pollet, Delcour, and Courtin 2010). The variances of activity exhibited by GH10 (1) against different substrates may be due to a combination of factors: the solubility of xylan and the amount and conservation of subsites present in the catalytic site structure of this enzyme. GH10 xylanases have a higher activity against soluble xylan fractions (such as quiona xylan and beechwood xylan) compared to a more insoluble substrate (*e.g*. birchwood xylan) (Linares-Pasten, Aronsson, and Karlsson 2018). In addition, the significant presence of X3 and X4 obtained as hydrolysis products from the evaluated xylans (**Figure 4**) would imply strong interactions of the substrate with −3, +3 and −4, +4 subsites of the active site of the enzyme, respectively. Conversely to substrate interactions more distal than −2 and +2 subsites may be weak or missing in previously described enzymes within this family (Salas-Veizaga et al. 2017).

In addition, the molecular model obtained for GH10 (1) indicates that some subsites close to the cleavage site of the enzyme do not allow substitutions in the xylose backbone chain, in particular the subsite −2. This subsite is highly interesting since arabinose-substitutions at this position can have a role in substrate recognition (Xie et al. 2006). In other GH10 family enzymes this position is conserved, and various studies have verified its ability to accept Araf substituent residues (Aronsson et al. 2018), so that these substitutions on the xylose backbone are not a big hindrance for arabinoxylan hydrolysis (Linares-Pasten, Aronsson, and Karlsson 2018). This could explain the low amount of X1 obtained from different xylans and the limited activity displayed on arabinoxylan (highly substituted, highly soluble).

To efficiently degrade arabinoxylan, PUL 15 from *P. copri* encodes an extracellular GH43_12. The substrates specificity of GH43_12 (**Table 5**) showed that the enzyme displays α-L-arabinofuranosidase activity (EC 3.2.1.55) and might be classified into type-B: enzymes active against small substrates, such as *p*-NP-Ara and arabinoxylooligosaccharides (AXOS). However, in addition to this, the enzyme is likely to catalyze the hydrolysis of arabinose from polymeric substrates, such as arabinoxylans (Yang et al. 2015). These characteristics would indicate that this enzyme acts synergistically with GH10 in substituted xylan, where GH43_12 removes arabinose decorations, thus enhancing the GH10 activity on arabinoxylan. Moreover, GH43_12 might act on AXOS generated by GH10 enzymes, debranching such oligosaccharides to be transported to intracellular location. There, those oligosaccharides act as substrates of the exoxylanase/xylosidase GH43_1. The separate action of GH10 and GH43_12 can also explain the accumulation of arabinose first seen in some of the growth trials of *P. copri*, indicating a faster use of nonsubstituted xylan, but which by extended cultivation allowed debranching and further use of the substrate.

GH43_12 (PUL 10) from *P. copri* DSM 18205 showed a low α-L-arabinanase activity (EC 3.2.1.99), contrary to other GH43 arabinofuranosidases (AFs) identified in probiotic microorganisms, such as LbAraf43 and WAraf4 from *Lactobacillus brevis* DSM1269 and *Weissella* strain 142, respectively (Linares-Pastén et al. 2017), which showed an almost exclusive specificity for 1,5-linked arabinooligosaccharides (AOS) while they were not active with rye arabinoxylans (a xylan polymer substituted by 1,3- or 1,2-linkages arabinofuranosyl branches). Arabinanase activities may however be covered by another PULs in *P. copri*, such as PUL11 that encodes two putative arabinanases homologous to candidates from GH43 subfamilies (4 and 5).

Finally, the intracellular GH43_1 (PUL 15) showed β-xylosidase, arabinofuranosidase and exo-1,4-β-xylanase activity (E.C. 3.2.1.37) according to the substrate specify results (**Table 5**). Exo-β-xylanases usually show multiple enzyme functions which are helpful for a more efficient degradation of xylan (Juturu and Wu 2014). The presence of GH43 enzymes that display exo-xylanase activity within *Prevotella* species was previously reported in *P. ruminicola* B_1_4 (**XynB**) (Gasparic et al. 1995). Similar to GH43_1 from *P. copri* DMS 18205, XynB exhibited activity against *p*-NP-Xyl, and *p*-NP-Ara, and was active against birchwood xylan. Both enzymes lack the native N-terminal signal peptide, therefore they are expected to be cytoplasmic proteins. The similarity between these two enzymes is also observed by BLAST analysis of the translated sequence of GH43_1, which showed an identity of 79.9% with XynB (**Table 1**), being the closest related sequence found in UniProtKB Swiss-Prot/PDB Database. The analysis of hydrolysis products obtained from different xylans would suggest the mode of action of

GH43_1. For xylans from birchwood, beechwood, and quinoa (**Figure 4**), the exo-β-xylanase continuously cleave xylose from the end of the xylan chain, until it is blocked by the substituent groups of these xylan types (MeGlcA residues). Similar results were reported with the XynB enzyme from *P. ruminicola* B_1_4 in the hydrolysis of xylan from oat spelt and birchwood, where aldotetrauronic acid substitutions inhibit enzymatic activity (Gasparic et al. 1995).

It has been reported that the substrate specificities of GH43 β-xylosidases present in microorganisms with probiotic characteristics and isolated from the GI tract, correlate with the preference of those bacteria to assimilate xylooligosaccharides. In this sense, the GH43 enzymes reported in *Lactobacillus brevis* DSM 20054 and *Weissella* sp. strain 92, now identified as *Weissella cibaria* strain 92 (Månberger et al. 2020), showed a higher activity against X2 and X3 than that on xylooligosaccharides of higher DP (Falck et al. 2016; Michlmayr et al. 2013). In addition, growth assays in a mixture of xylooligomers of different DP showed that these microorganisms assimilated X2 and X3 faster than xylooligosaccharides of higher DP (X4, X5, and X6), showing a clear preference for short-chain substrates. (Mathew et al. 2018; Patel et al. 2013). In probiotic microorganisms with an outstanding capacity to assimilate and grow in the presence of xylooligosaccharides of higher DP, such as *Bifidobacterium animalis* subsp. *lactis* BB-12, a GH43 β-xylosidase (Viborg et al. 2013) was identified that presented a specific activity for X2 2.3-fold higher than for *p*NP-Xyl and increased even more for X3 (3.6-fold higher) and X4 (5.6-fold higher). In *B. adolescentis* ATCC 15703 a GH120 β-xylosidase (xylB) was identified, which shows a drastic increase in the specific activity from X2 to X4 (Lagaert et al. 2011) and, remarkably, this microorganism achieved the total consumption of a mixture of xyloligosaccharides (X2, X3, X4, and X5) after a 48 h fermentation (Falck et al. 2013).

In these microorganisms it can be seen how, through evolutionary adaptation in a competitive and constantly changing environment such as the GI tract, the different xylanases present various specificities and catalytic efficiencies for the utilization of oligosaccharides that other species cannot assimilate. *P. copri* has been shown to be associated to a diet related to dietary fibers incorporation, where the selective pressure for the use of polysaccharide complexes is greater than other diets (Kovatcheva-Datchary et al. 2015; Tett et al. 2019). Multi-omics studies have also confirmed that AXOS intake increased the proportion of *Prevotella* species and particularly *Prevotella copri* among the bacterial species of the analyzed fecal microbiome (Benítez-Páez et al. 2019). In this sense, because the enzyme GH43_1 from PUL 15 of *P. copri* DSM 18205 presents a catalytic efficiency for X5 superior to the other tested substrates (although with the fastest kcat on X2 and X3), we hypothesize that this strain presents the ability to take up and efficiently degrade oligosaccharides with a higher DP than those reported for *Lactobacillus* species, an advantage also previously studied in probiotic *Bifidobacterium* species (Falck et al. 2013; Lagaert et al. 2011; Viborg et al. 2013). Additional studies of xylooligosaccharides fermentation and uptake are however necessary to confirm this hypothesis.

Based on the results and analyzes in this work, we propose the following model of xylan degradation by the synergistic action of *P. copri* DMS18205 enzymes (**Figure 8**). Extracellular α-L-arabinofuranosidase type-B, GH43_12 from PUL 10, is capable of debranching arabinoxylans (**Table 5**) generating a poorly branched xylan which in turns act as a substrate for the extracellular GH10 (1) endo-β-xylanase from PUL 15. This enzyme hydrolyzes the β-1,4 glycosidic bonds of the xylan backbone, and due to the presence of many catalytic subsites in the enzyme (**Figure 7C** and **Table 5**), the final products are mainly xylooligosaccharides with a high DP (**Figure 4**). Other microorganisms of the GI tract would not assimilate these high-molecular-weight sugars; instead, they would incorporate into the cellular interior of *P. copri*. These products are substrates of intracellular β-xylosidase with exo-xylanase activity GH43_1 from PUL 15, which has activity against oligosaccharides in a DP-range from 2-5 (**Table 4**), generating xylose as the main hydrolysis product (**Figure 4**) that would be incorporated into the classical routes of assimilation of pentoses. This synergistic and specific interaction of the three described enzymes from *P. copri* DSM 18205 might imply a competitive advantage developed by this microorganism to utilize xylan fraction of diary fibers in the complex environment of the mammalian gut.

**Figure 8.**
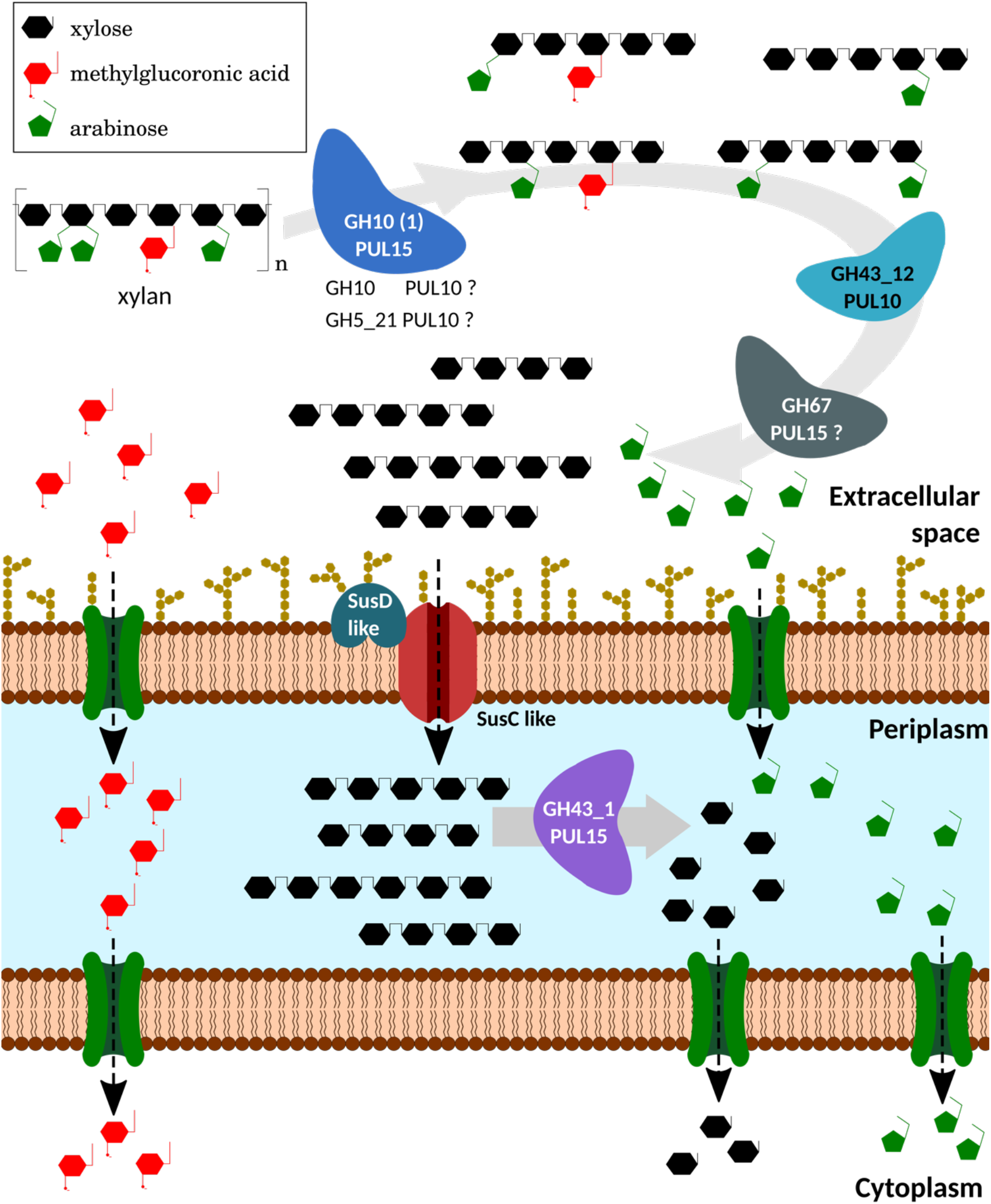
Depolymerization model of xylan by *Prevotella copri* DMS18205, based on genomic and biochemical studies of its produced enzymes. In the extracellular space, the endo-1,4-β-xylanase GH10 (1) from PUL 15 (and, possibly, GH10 and GH5_21 from PUL 10) would act, decreasing the degree of polymerization of xylan. The reaction products, xylooligosaccharides substituted with a high DP, can be debranched by the action of extracellular α-arabinofuranosidase GH43_12 from PUL 10 (and, according to genomic studies, by α -glucuronidase GH67 from PUL 15). These oligosaccharides would be introduced into the cell using a variety of proteins like SusC and SusD. In the periplasmatic region, the xylooligomers obtained would be degraded by β-xylosidases of the families GH43_1 from PUL 15. In this way, monomers that can enter common metabolic pathways would be produced. (Illustration elaborated using vectorial graphic editor *Inkscape v1.0*)

## Conclusions

Two PULs in *Prevotella copri* DMS18205 were described as potential candidates involved in xylan degradation, from which three novel enzymes were identified, produced, and characterized.

PUL10 encoded a GH43, subfamily 12 enzyme (GH43_12) with α-arabinofuranosidase activity of type-B with specificity against arabinoxylans, allowing debranching of single substituted arabinose groups and with the ability to hydrolyze arabinose substituents. PUL15 presented two genes encoding an endo-β-xylanase from GH10 and an exo-xylanase/ beta-xylosidase, from GH43, subfamily 1 (GH43_1). The GH10 enzyme was able to hydrolyse substituted xylans and presented a production profile of xylooligosaccharides with equal amounts of XOS in the DP-range 2-4, possibly due to the high number of identified catalytic subsites. The GH43_1, despite its probable intracellular localization, exhibited exo-β-xylanase activity with high affinity and efficiency to degrade higher DP oligosaccharides, which could mean an evolutionary advantage for the assimilation of XOS in the GI microbiome.

Although previous studies have demonstrated the assimilation of polysaccharides by *P. copri*, this work constitutes the first production and characterization of carbohydrate-active enzymes, essential to better understand the process of assimilation of xylan-derived polysaccharides by this microorganism.

## Acknowledgments

This work was supported by the Antidiabetic Food Center, funded by Vinnova, the ScanOats programme funded by SSF, the foundation Gyllenstiernska Krapperupsstiftelsen and the National Agency for the Scientific and Technological Promotion, Argentine, project ref. PICT-2017-2185, and BecAr Program, Argentine, for transfer and maintenance of J.S.H and J.H.P.

## Supplementary information

**S1.**
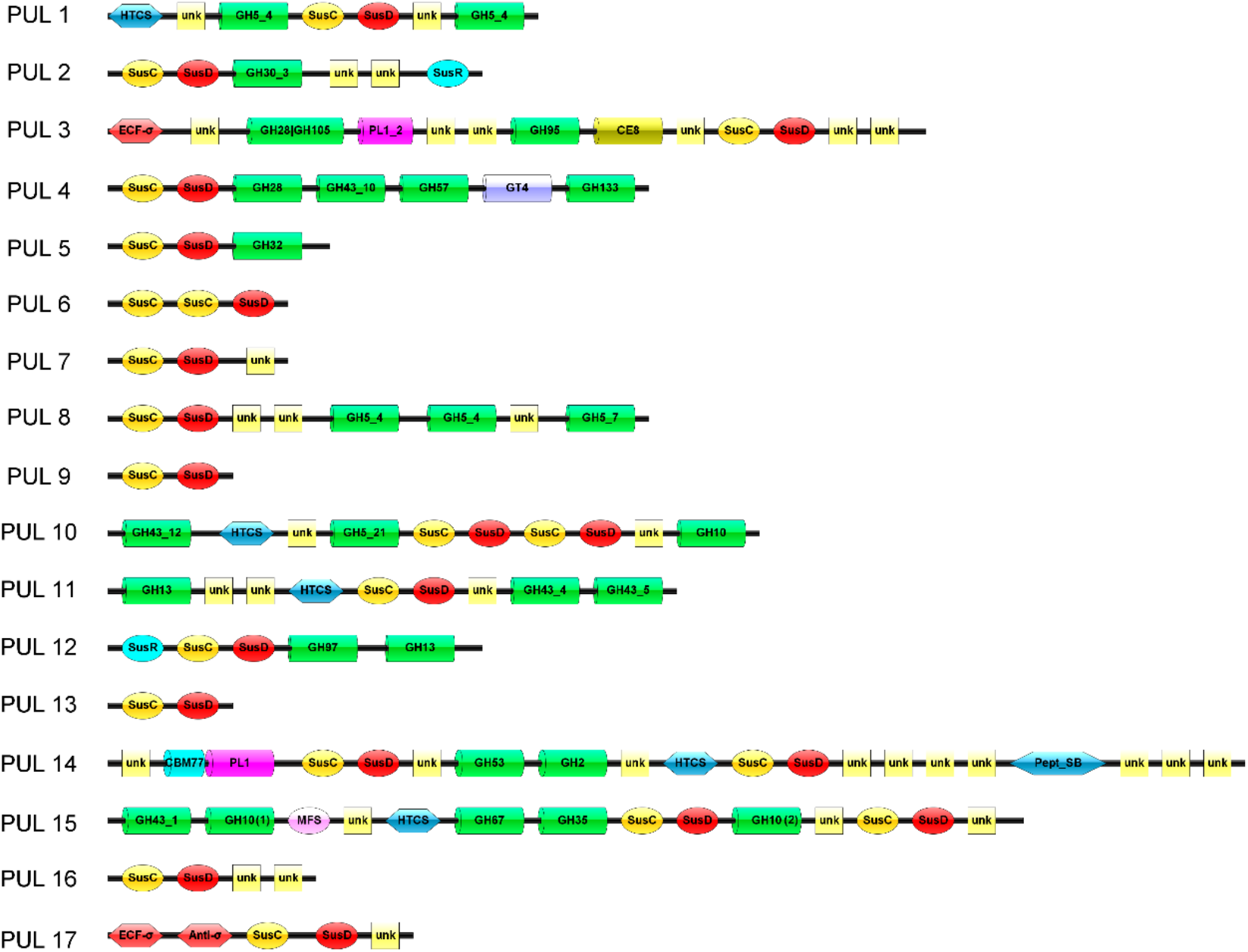
Modularity of the Polysaccharide Utilization Loci (PULs) predicted by the Polysaccharide-Utilization Loci DataBase (PULB) of *Prevotella copri* DMS 18205. **HTCS**: Hybrid Two-Component Systems sensor-regulator. **unk**: hypothetical protein. **GH**: Glycoside Hydrolase Family. **Sus**: Starch-Utilization System. **ECF-σ**: Extracytoplasmic Function Sigma Factor. **PL**: Polysaccharide Lyase Family. **CE**: Carbohydrate Esterase Family. **GT**: Glycosyl Transferase Family. **CBM**: Carbohydrate-Binding Module Family. **Pept_SB**: Peptidases Serine Family. **Anti-σ**: Extracytoplasmic Function Anti-Sigma Factor. **MFS**: Major Facilitator Superfamily permease.

**S2.**
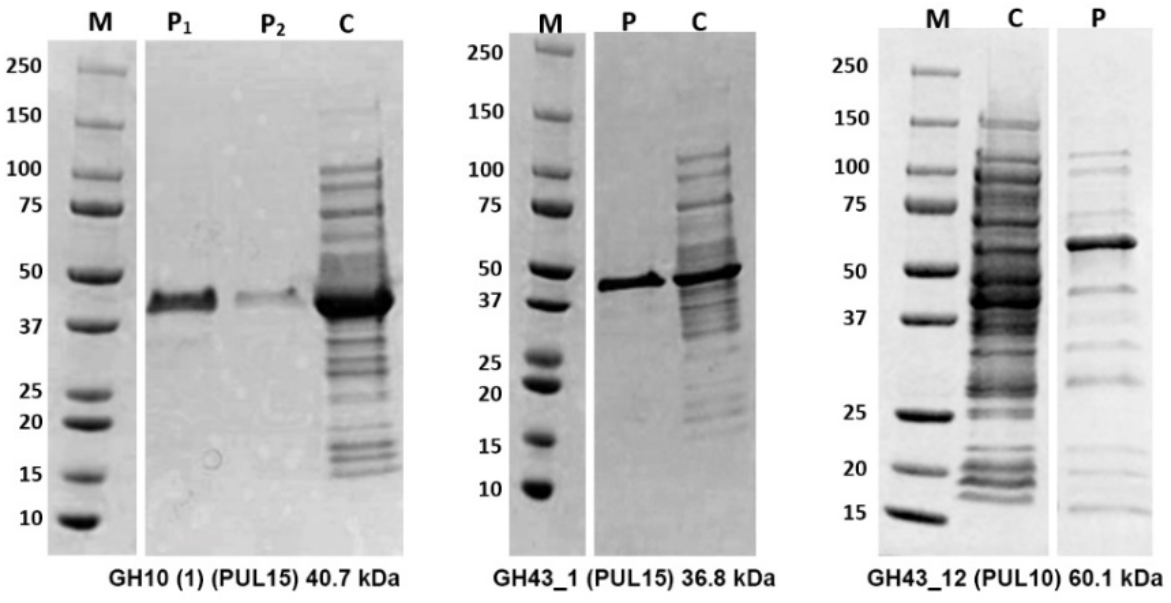
SDS-PAGEs showing GH10 (1), GH43_1 and GH43_12 proteins from *Prevotella copri* DSM 18205 overexpressed and purified. Molecular weight marker in kDa (**M**), crude extract (**C**), and the purified protein from the soluble fraction (**P**).

**S3.**
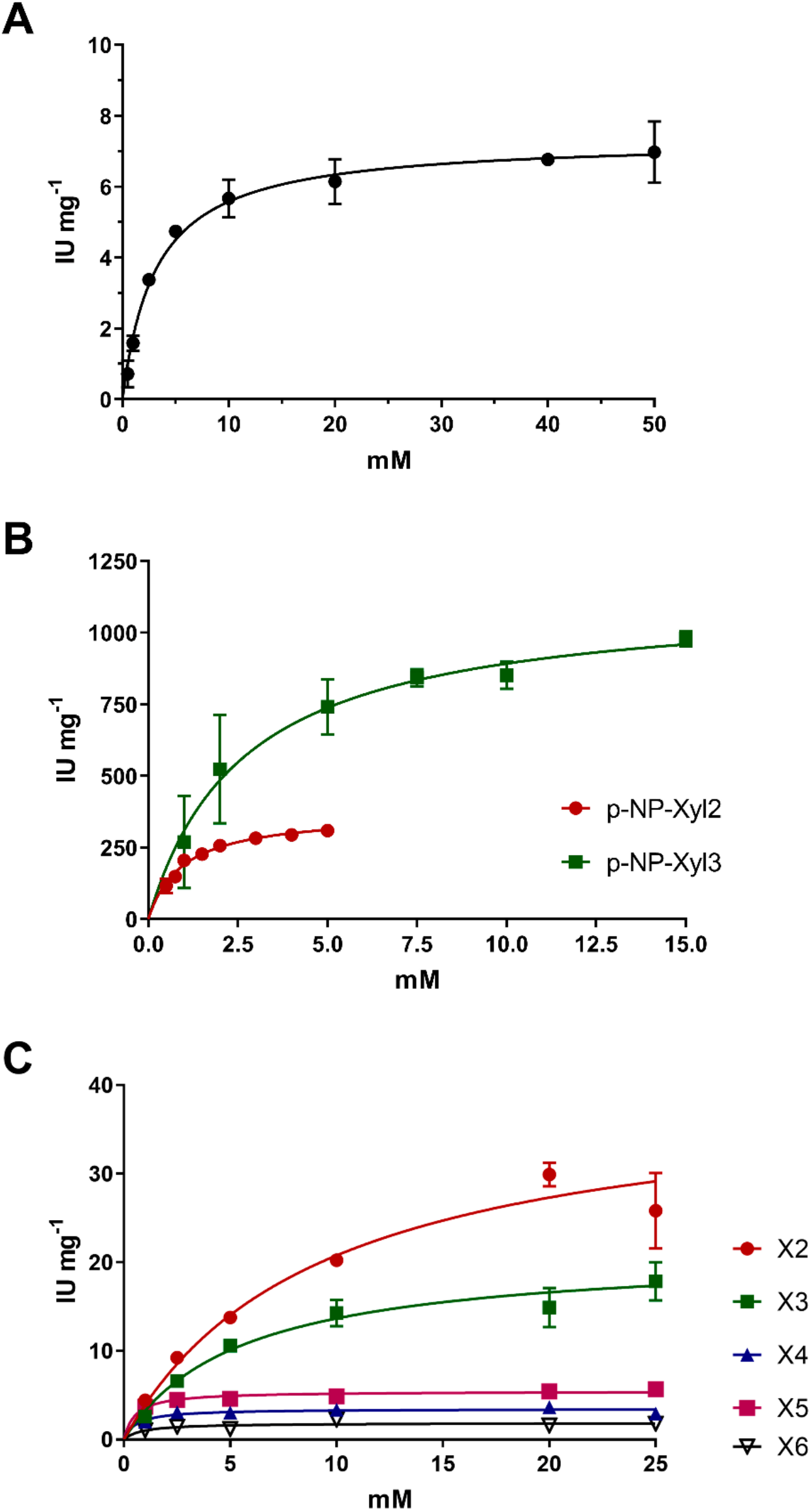
Nonlinear regressions to the Michaelis–Menten equation at 37°C and pH 5.5. Error bars represent the standard deviation for n =3. (**A**) GH43_12 (PUL 10), substrate: p-NP-α-L-arabinofuranoside. (**B**) GH10 (1) (PUL 15), substrates: p-NP-Xyl2 (p-Nitrophenyl-β-xylobioside) and p-NP-Xyl3 (p-Nitrophenyl-β-xylotrioside). (**C**) GH43_1 (PUL 15), substrates: X2 (xylobiose); X3 (xylotriose); X4 (xylotetraose), X5 (xylopentaose) and X6 (xylohexaose).

**S4.**
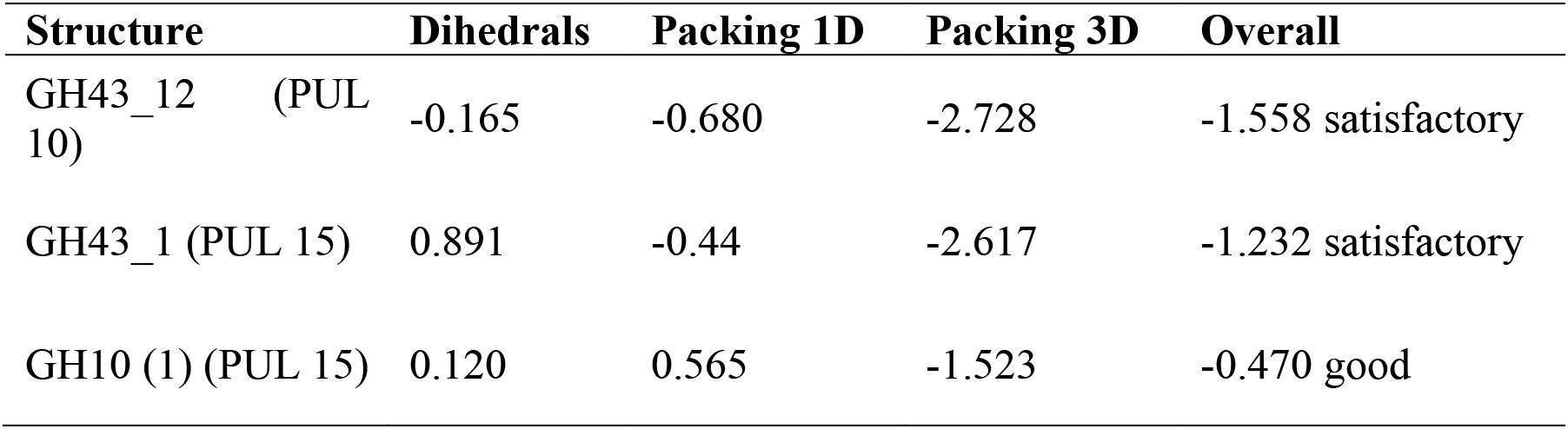
Molecular models validation. Quality Z-scores of the hybrid models obtained using YASARA.

**S5.**
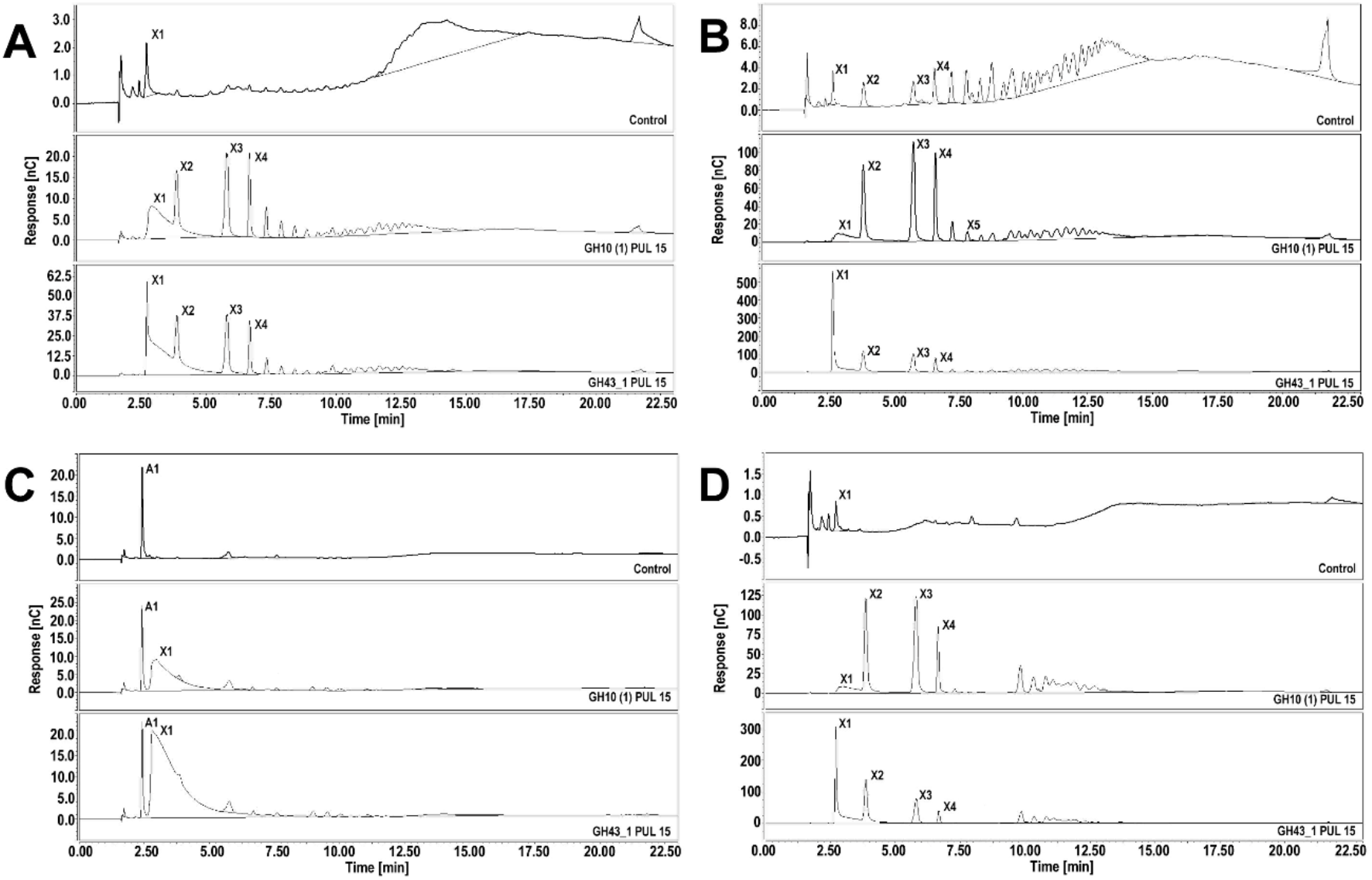
Chromatograms ofhydrolysis products from enzymatic degradation using the PUL15 enzymes from GH10 and GH43_1, along with substrate controls. Notice beechwood xylan consisted of oligomers which affected the yield of the enzymatic hydrolysis. Birchwood xylan (**A**), beechwood xylan (**B**), arabinoxylan (**C**) and quinoa xylan (**D**). X1, xylose; X2, xylobiose; X3, xylotriose; X4, xylotetarose; X5, xylopentaose.

